# Supramolecular Site-Specific Antibody Drug Conjugates Outperform Cysteine-Conjugated Analogs in Pancreatic Peritoneal Carcinomatosis

**DOI:** 10.64898/2026.08.05.743073

**Authors:** Christopher M. Gromisch, Alina Ringaci, Aladin Hamoud, Samantha M. Berry, Mariana Gromisch, W. Mark Saltzman, Mark W. Grinstaff

**Author notes:** Corresponding authors: Christopher M. Gromisch, Room 401E, 55 Prospect Street, New Haven CT New Haven, CT 06511, Mark W. Grinstaff, Room 519, 590 Commonwealth Ave, Boston MA, Boston, MA 02215.

## Abstract

Antibody drug conjugates (ADCs) are a burgeoning class of targeted therapies. However, limitations in their synthesis and efficiency of payload delivery restrict their clinical utility. Here we report a supramolecular assembly (SMA) ADC conjugation method, which allows site-specific, uniform drug loading, resulting in an enhanced pharmacokinetic profile and *in vivo* efficacy. This peptide conjugation strategy relies on spontaneous heterotetrameric coiled-coil formation between a pair of peptides appended on the C-terminus and a drug-loaded complementary pair in aqueous solution. Pairing this SMA conjugation with an antibody that targets the dual-endothlin-1/VEGF signal peptide receptor (DEspR), a pancreatic ductal adenocarcinoma (PDAC) specific receptor, retains antibody binding and plasma stability. When the anti-DEspR monoclonal antibody is conjugated with monomethyl auristatin E (MMAE), the ensuing ADC internalizes following cell surface binding and induces selected cell death in multiple DEspR positive PDAC cell lines. *In vivo*, the ADC exhibits favorable pharmacokinetics, high tumor specificity, and improves overall survival in a rat orthotopic model of pancreatic peritoneal carcinomatosis, compared to conventional ADC conjugation. A heterotetrameric coiled-coil structure enables the efficient synthesis of a potent ADC, further documenting the versatility of supramolecular scaffolds as key orthogonal building block for site-specific conjugation in biopharmaceutical and biomaterial drug delivery systems.

**One Sentence Summary:** Combining site-specific conjugation of monomethyl auristatin E, via the use of biologically inspired heterotetrameric coiled-coils, with a tumor-selective antibody targeting the dual-endothlin-1/VEGF signal peptide receptor affords a highly effective, ADC, which improves survival in a rat orthotopic model of pancreatic peritoneal carcinomatosis compared to standard cysteine-conjugated analogues with higher drug loading.

## INTRODUCTION

Antibody-drug conjugates (ADCs) are a rapidly evolving class of directed cancer therapies and combination therapeutics. The concept of ADCs originated in the 1960s(*1*); however, the first FDA approved ADC, gemtuzumab ozogamicin (Mylotarg^TM^), was not commercially available until the early 2000s.(*2*) Today, fifteen ADC have received regulatory approval for the treatment of breast cancer, leukemia, lymphoma, ovarian cancer, cervical cancer, multiple myeloma, and bladder cancer.(*3*) Advances in drug discovery, linker and conjugation chemistry, and antibody target identification are fueling progress, but the vast majority of US ADC clinical trials are no longer being pursued.(*3*) The shortcomings of ADCs include poor target selection and unfavorable pharmacokinetics and pharmacodynamics from drug conjugation.(*4*) Therefore, there is considerable interest in improving ADC target selection and design to ensure safe, effective drug delivery.

Effective ADC design requires careful consideration of the treatment target and method of drug conjugation. Ineffective drug conjugation significantly hampers ADC performance and results in high variability in drug-to-antibody ratios (DARs), significant product heterogeneity, increased aggregation, altered antibody binding, increased toxicity, and decreased circulating ADC half-life.(*5,6*) Although advances have been made in conjugation strategies, most FDA-approved ADCs conjugate exposed lysine or reduced cysteine residues.(*4,5*) Current strategies for site-specific drug conjugation, which include enzymatic conjugation, terminal sequence tags, unnatural amino acids, and engineered natural amino acid sequences (see Table S1) are improving ADC pre-clinical performance.(*7,8*) However, even with these advances, limitations still persist in: efficacy and consistency of conjugation, reductions in antibody stability from the use of denaturing solvents, and antigenicity of ADC design. Overcoming these limitations and enhancing the consistency of conjugation and preparation of uniform drug products should further improve ADC performance and advance new targeted therapies.

Given our clinical interest in pancreatic cancer and the dearth of available treatments (*9–12*), we report a novel ADC that targets the dual-endothlin-1/VEGF signal peptide receptor (DEspR), which is specific to pancreatic ductal adenocarcinoma (PDAC) (Fig. 1A). DEspR is highly expressed on human PDAC tumors, and regulates tumor vasculoangiogenesis, anoikis resistance, stress survival, and tumor invasion.(*13,14*) DEspR inhibition reduces tumor collagen expression,(*13,14*) which is of particular interest in ADC development, as this could improve drug penetrance in a desmoplastic tumor environment.(*15,16*) We enhance anti-DEspR ADC design through site-specific conjugation using a supramolecular(*17–21*) assembly (SMA) methodology (Fig. 1B). Specifically, a high affinity heterotetrameric coiled-coil structure forms between pairs of optimized leucine zipper peptides. Recombinant technology installs one set of coils onto the C-terminus of the antibody, while the other set, prepared by solid-phase synthesis, contains the drug for subsequent conjugation by self-assembly. Through this SMA method, we prepare ADCs with uniform loading of two monomethyl auristatin E (MMAE) per antibody, under mild aqueous conditions, without impacting antigen binding. We find that the SMA ADC, α-DEspR_SMA_-MMAE_2_, demonstrates pro-longed *in vivo* circulation and more favorable pharmacokinetics than a traditional cysteine conjugated product, and improves survival and reduces tumor burden in a rat xenograft model of pancreatic cancer peritoneal carcinomatosis.

**Fig. 1.**
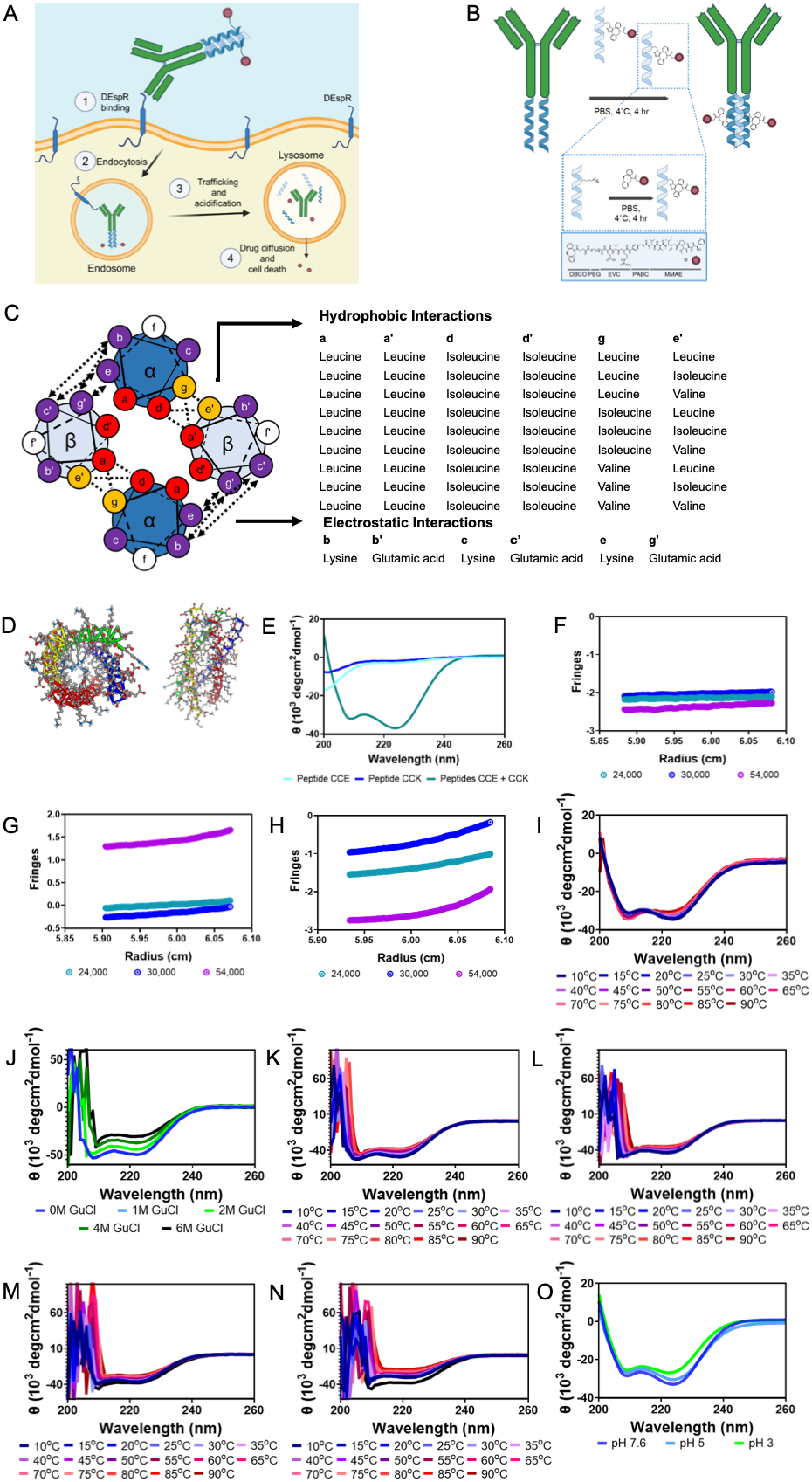
Biophysical quantification of coiled coil design. **(A)** Overview of ADC internalization: ADCs bind cell-surface DEspR, are trafficked to the lysosome, and release free MMAE through cathepsin-B cleavage of the linker. (**B)** Overview of SMA conjugation: conjugation occurs first on the CCE-peptide, which is purified, and then reacts under mild, aqueous conditions with the recombinantly modified α-DEspR-CCK. **(C)** Wheel diagram demonstrating the tested amino acids and configuration of experimental peptides. **D)** Three-dimension modeling of coiled-coil structure of CCE-CCK interaction using CC Builder 2.0. (**E)** CD spectrum of CCE (teal) and CCK (blue), and CCE-CCK (green). (**F-H)** Analytical ultracentrifugation equilibrium analysis of (**F**) peptide CCE (single species), (**G)** CCK (single species) and (**H)** CCE-CCK (tetramer, K_d_=1.1021×10^-10^). (**I)** Thermal denaturing CD spectrum of CCE-CCK, with 18.2±0.8% denaturation at 90^°^C. (**J)** Chaotropic denaturing CD spectrum of CCE-CCK, with 25.7±0.9% denaturation at 6M guanidium chloride (GuCl). **(K-N)** Combined chaotropic-thermal denaturing CD spectrum of CCE-CCK at (**K)** 1M, (**L)** 2M, (**M)** 4M, and (**N)** 6M GuCl. CCE-CCK was 53.5±0.8% unfolded in 6M GuCl at 90^°^C. **O)** CD spectrum of CCE-CCK at pH of 3 (green), 5 (light blue), or 7.6 (dark blue). CCE-CCK helical structure was reduced by 7.3% and 16.9% in pH 5 and pH 3 solutions.

## RESULTS

### Designed peptides form a high-affinity, high stability heterotetramer

We designed a series of peptides to form parallel heterotetrametric coiled-coils based on the work by Harbury et al(*22*) (Fig. 1C, peptide sets represented as either blue or light blue). To favor a single oligomeric state, we fixed the *a* position in the heptad as leucine and the *d* position as isoleucine (Fig. 1C, red). We then modified the *g* or *e’* of each peptide to either have β-branched (lysine and valine) or β-unbranched (isoleucine) amino acids, to increase van-der-Waals interaction and improve coil strength(*23*) (Fig. 1C, orange). To minimize self-interaction, we designed the *b*/*c* positions to be positively charged (lysine residues) and the *b’*/*c’* positions to be negatively charged (glutamic acid, Fig. 1C, purple; see Table S2 for the full structures). We selectively modified a single *f’* residue to include azidolysine for future dicyclobenzyloctene drug conjugation (Fig. 1C, white). The overall predicted structures, using CCBuilder 2.0(*24*), form a complementary tetrameric package with a suprahelical diameter approximating the distance between antibody heavy chains (Fig. 1D). We screened each species using circular dichroism to assess the individual peptide structure and equimolar solutions were analyzed to determine both the formation and structure of heterooligomeric species. (Fig. S1A-O). The substitution of valine at the *e’* (CCE-peptide) and *g* (CCK-peptide), fit this criterion, with a left-handed coiled-coil in the heteromeric state (ratio of 222/208 nm>1.1) (Fig. 1E), and helical character >99%.(*25*) Individually, the CCE and CCK demonstrate disorganized structures.

We used analytical ultracentrifugation to confirm heterotetrameric coiled-coil formation through velocity scans of CCE, CCK, and CCE-CCK. CCE and CCK form discrete low-molecular weight species (estimated molecular weight <4,000 kDa, Fig. S2A,B), while CCE-CCK gives a predominant single species with a molecular weight of 16,281 kDa in accordance with predicted molecular weight of heterotetrameric coiled-coil structure (theoretical 15,788 kDa) (Fig. S2C). We confirmed the above findings with equilibration analysis using three rotation speeds based on theoretical sedimentation, 24,000, 30,000, and 54,000 rpm, along with three concentrations for peptide CCE, CCK, and CCE-CCK: 0.2, 0.6, and 1.0 mg/mL (Fig. 1F-H, Fig. S3A-C). Experimentally determined species buoyant molecular weights (M _B_) along with the root mean square deviation (RMSD) of the equilibration data and global score chi-square (GBCS) are shown in Table 1.

**Table 1.** Single Species Analysis of Peptides CCE, CCK, and CCE-CCK.

| Species | $M_B$ ($M_w$) | RMSD | GBCS |
| --- | --- | --- | --- |
| Peptide CCE | 4241 (3990) | 0.005422 | 1.175805 |
| Peptide CCK | 4288 (3904) | 0.005684 | 1.1292114 |
| Peptide CCE-CCK | 15306.316 (15788) | 0.005791 | 1.213398 |

We next modeled the species as monomers, dimers, trimers, and tetramer aggregates with mass conservation, with model parameters based on optimization from monomer analysis. The RMSD of fit for each peptide in different oligomeric states is provided in Table S3 and the best fit data and prediction dissociation constants are provided in Table S4. Individual peptides CCE and CCK best fit a single-species model, with weak dissociation constants noted for a monomer/tetramer self-association model (K_d_,_CCE_: 0.995 M and K_d_,_CCK_: 0.0971 M). Equimolar CCE-CCK best fit a tetrameric-species, with a K_d,(CCE-CCK)(CCE-CCK)_=1.1021×10^-10^. Together, these data suggest that CCE-CCK formed a highly stable tetramer.

Next, we used circular dichroism with thermal and chaotropic denaturation, providing biophysical assessment of peptide affinity, structure reversibility, and aggregation risks.(*26,27*) Thermal denaturing reveals minimal helical change (18.2±0.8% unfolding at 90^°^C), with complete reversibility and maintenance of helical character after sequential thermal runs (n=3) (Fig. 1I, Fig S4A). Chaotropic denaturation using guanidium chloride (GuCl), reduces coil stability by 25.7±0.9% at 6M GuCl; however, helical structure is maintained (Fig. 1J, Fig. S4B). With combined chaotrope and thermal denaturation, the coiled coil structure denatures by 53.5±0.8% (Fig. 1K-N). Collectively, these data suggest that the CCE-CCK interaction is of high affinity and stability, and appropriate as a non-covalent linkage for ADC design.

Finally, we characterized the impact of pH and salt on coiled-coil stability, given the importance of salt bridge interactions (Fig. 1C). Addition of hydrochloric acid to phosphate buffered saline (mimicking movement through sorting endosomes to lysosomes), does not significantly perturb coiled-coil stability at pH 5 (7.3% loss of helical character) or pH 3 (16.9% loss of helical character) (Fig. 1O). Furthermore, thermal denaturing experiments at pH 3 show similar reversibility as observed at pH 7.6, suggesting that changes in salt bridge do not adversely affect the coiled-coil stability (Fig. S4C). The formation of a coiled-coil structure depends on salt-bridges, as a sodium chloride between 125-250 mM is needed for CCE-CCK formation (Fig. S4D). Reducing the salt concentration collapses the coiled structure into structurally disorganized peptides (Fig. S4D).

### Supramolecular assembly conjugation enables ADC preparation

We recombinantly inserted the CCK-peptide with a GGGGS linker into the C-terminus of each heavy chain of a dual-endothlin-1/VEGF signal peptide receptor (DEspR) monoclonal antibody (α-DEspR-CCK mAb), based on the previously reported 7c5 antibody.(*14*) The C-terminal CCK allows for SMA conjugation once mixed with the drug-loaded CCE-peptides (Fig. 1B). A reducing SDS-PAGE confirms the addition of CCK (Fig. S5A). Next, we chemically grafted CCE with either DBCO-PEG_4_-valine-citrulline-PABC-MMAE, DBCO-PEG_4_-glutamic-acid-valine-citrulline-PABC-MMAE, DBCO-AF488, or DBCO-AF647, and confirmed loading by RP-HPLC (Fig. S5B-I). We first synthesized ADCs using DBCO-AF647 or a DBCO-PEG_4_-glutamic-acid-valine-citrulline-PABC-MMAE linker grafted onto CCE mixed in a 2.1:1 ratio with α-DEspR-CCK (Fig. 1B). Solution addition of the CCE to α-DEspR-CCK forms a single species of either a conjugate with two AF647 fluorophores (α-DEspR_SMA_-647_2_, Fig. 2A) or two MMAE molecules (α-DEspR_SMA_-MMAE_2_, Fig. 2B), evaluated by non-reductive RP-HPLC. By UV-VIS, the calculated DAR is 2.00±0.14 and 2.08±0.15 for α-DEspR_SMA_-647_2_ and α-DEspR_SMA_-MMAE_2_, respectively. Coupling of DBCO-AF488 and DBCO-PEG_4_-valine-citrulline-PABC-MMAE to α-DEspR-CCK affords similar values (2.06±0.10 and 2.10±0.13, respectively; Fig S6A-G). To confirm the selectivity of this conjugation method, we performed non-reductive RP-HPLC of α-DEspR-CCK mixed with CCE-DBCO-AF647 compared to α-DEspR mAb mixed with CCE-DBCO-AF647. The presence of the C-terminal CCK allows conjugation of the AF647 (Fig. 2C, top blue lines), while the absence of it results in two separate peaks (Fig. 2C, middle teal lines) of the antibody and CCE-DBCO-AF647. We similarly evaluated if this selectivity extends to the DBCO-PEG_4_-glutamic-acid-valine-citrulline-PABC-MMAE linker, which is more hydrophobic and could interact with hydrophobic pockets on α-DEspR mAb. Similarly, the DBCO-PEG_4_-glutamic-acid-valine-citrulline-PABC-MMAE assembles with α-DEspR-CCK (Fig. 2D, top blue lines), but not with α-DEspR mAb (Fig. 2D, middle teal lines).

**Fig. 2.**
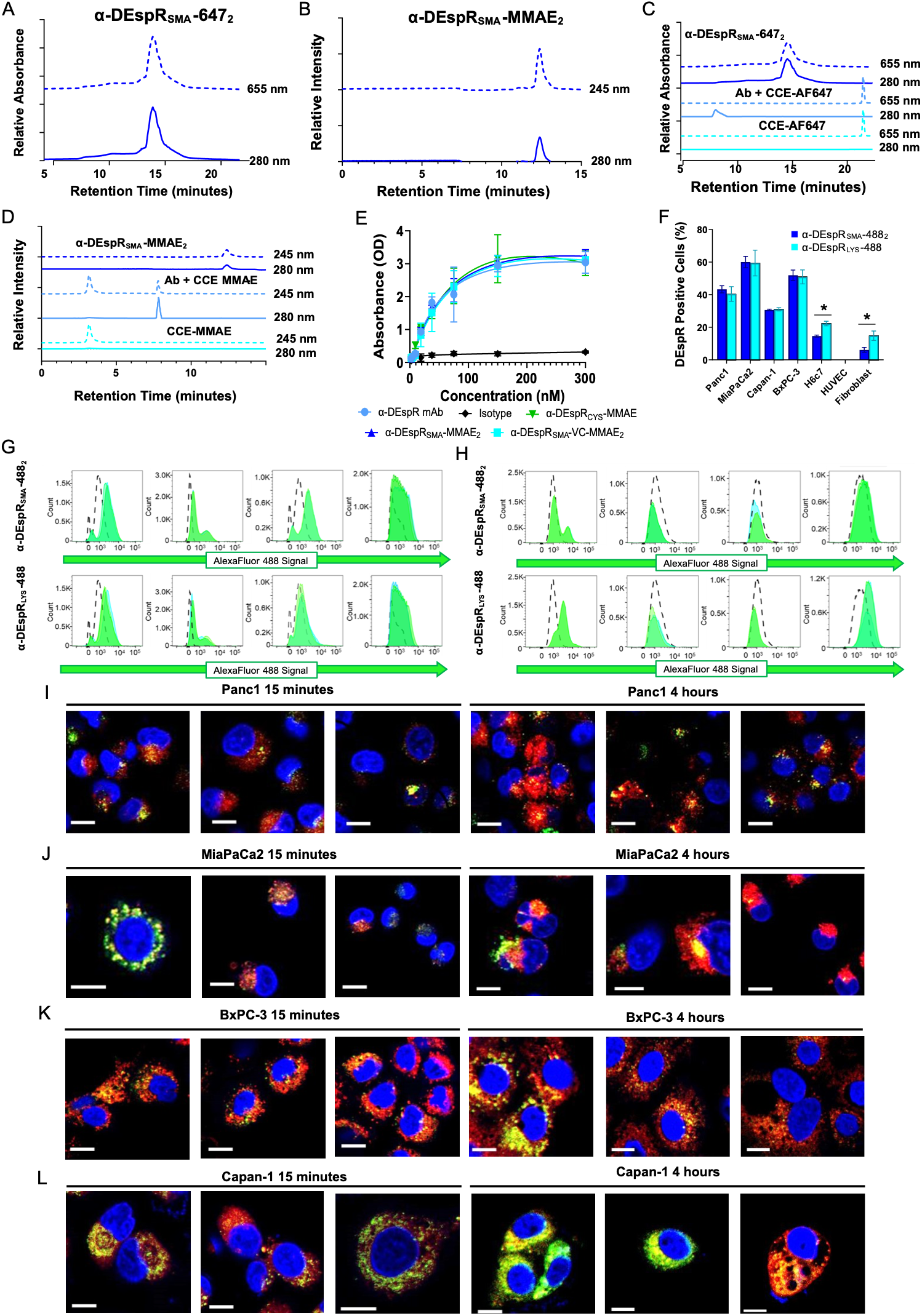
**ADC Synthesis and Binding Characterization**. **(A,B)** Representative RP-HPLC of α-DEspR-CCK conjugated with CCE-AF647 (CCE+DBCO-AF647) and CCE-EVC-MMAE (CCE+DBCO-PEG_4_-glutamic acid-valine-citrulline-PABC-MMAE), showing single products α-DEspR_SMA_-647_2_ and α-DEspR_SMA_-MMAE_2._ **(C,D)** Representative RP-HPLC showing the specificity of SMA conjugation in α-DEspR_SMA_-647_2_ and α-DEspR_SMA_-MMAE_2_ synthesis. The top (blue) chromatograms show a 2:1 mixture of CCE-AF647 or CCE-EVC-MMAE with α-DEspR-CCK, while the middle (teal) show a 2:1 mixture of CCE-AF647 or CCE-EVC-MMAE with α-DEspR mAb, demonstrating that ADC synthesis requires CCE-CCK interaction. **E)** Binding of α-DEspR mAb (black, K_d_=52.93±11.23 nM), IgG2b isotype (purple), α-DEspR_CYS_-MMAE (blue, K_d_=51.45±10.12 nM), α-DEspR_SMA_-MMAE_2_ (red, K_d_=55.24±8.195 nM), and α-DEspR_SMA_-VC-MMAE_2_ (orange, K_d_=51.24±9.60 nM) to antigenic peptide M_1_TMFKGSNE_9._ (**F-H)** Binding of cell-surface DEspR using **(F)** lysine-based conjugation (α-DEspR_LYS_-488, light blue) and SMA conjugation (α-DEspR_SMA_-488_2,_ blue). Total DEspR-positive populations were identical in (**G)** PDAC cell lines, though DEspR_SMA_-488_2_ allowed better discrimination of low/high DEspR populations, while α-DEspR_LYS_-488 had higher staining in **(H)** DEspR negative/low control cells. (* p< 0.05) **(I-L)** Representative confocal microscopy showing internalization of α-DEspR_SMA_-647_2_ (red) in PDAC cell lines at 15 minutes vs. 4 hours. Cellular stains were NucBlue (blue, nuclei) and Lysotracker (green, lysosome). Colocalization of α-DEspR_SMA_-647_2_ with the lysosome (yellow-orange), increased over time in all PDAC cell lines, supporting use of lysosomally cleaved linkers for drug delivery.

We next characterized *ex vivo* drug release and peptide dissociation from α-DEspR_SMA_-MMAE_2_. We monitored ADC samples (0.1 mg/mL) in PBS or rat serum at 37°C over 30 days for MMAE and peptide dissociation by RP-HPLC. In PBS, the ADCs aggregate after 72 hrs without additives; however, there was no free CCE-MMAE or MMAE detected, within the limit of the instrument (Fig. S6H). Aggregation does not occur in serum over 30 days, and we detected no free CCE-MMAE and only trace amount of MMAE (2.9±1.5% of total free MMAE) after 30 days incubation (Fig. S6H).

### Supramolecular assembly conjugation does not impact antibody binding

We first tested whether SMA conjugation impacts antibody antigen recognition. We compared binding recognition to the target antigenic peptide (M_1_TMFKGSNE_9_) between α-DEspR mAb, SMA-conjugates α-DEspR_SMA_-VC-MMAE_2_ and α-DEspR_SMA_-MMAE_2_, and a commercially equivalent cysteine conjugate, using purchased maleimide-valine-citrulline-PABC-MMAE, assembled via disulfide reduction (α-DEspR_CYS_-MMAE) (*28*) (Fig S5G-I for characterization). The binding affinity is not statistically different between the native antibody (K_d_ 52.93±11.23 nM) and α-DEspR_SMA_-VC-MMAE_2_ (K_d_ 51.24±9.6 nM), α-DEspR_SMA_-MMAE_2_ (K_d_ 55.24±8.195 nM), or α-DEspR_CYS_-MMAE (K_d_ 51.45±10.12 nM). (Fig. 2E). We next compared binding of the SMA-conjugates to conventional labeling on DEspR-positive PDAC and DEspR-positive and negative control cell lines using flow cytometry. DEspR staining is equivalent between conventionally labeled (α-DEspR_LYS_-488) and SMA-labeled antibodies (α-DEspR_SMA_-488_2_) in PDAC cells, but conventionally labeled α-DEspR_LYS_-488 stains more DEspR-negative control cells (Fig. 2F). Furthermore, SMA labeled antibodies provide better discrimination of receptor-low and high populations than conventional labeling (e.g., MiaPaCa2: α-DEspR_SMA_-488_2:_ 27.3±2.6% vs. α-DEspR_LYS_-488 23.9±8.9%). (Fig. 2G,H).

Next, we performed confocal microscopy to study ADC internalization and trafficking. We used α-DEspR_SMA_-647_2_ as a surrogate to evaluate the effect of C-terminal modification on internalization compared to prior reported data.^14^ The α-DEspR_SMA_-647_2_ conjugate internalizes followed DEspR binding, with most α-DEspR_SMA_-647_2_ present near the cell membrane or adjacent to lysosomes at 15 minutes. Over time, α-DEspR_SMA_-647_2_ moves through the cell, with significant colocalization in the lysosome by 1 hour, up to a maximum of around 4 hours (Fig. 2I-L). We next evaluated internalization in DEspR-low (H6c7) and DEspR-negative (KV2) cells. Minimal uptake of α-DEspR_SMA_-647_2_ occurs in H6c7 cells, with only a few cells showing colocalization with the lysosome at 4 hours (Fig. S6I). Detectable uptake of conjugate is not seen in the DEspR-negative KV2, supporting the above data that SMA conjugation preserved receptor specificity (Fig. S6J)

### SMA bioconjugates α-DEspR_SMA_-VC-MMAE_2_ and α-DEspR_SMA_-MMAE_2_ enhance antibody efficacy in PDAC cells

We characterized whether SMA-generated ADCs effectively target PDAC cells, while maintaining safety in DEspR-negative cells. We selected 3-(4,5-dimethylthiazol-2-yl)-5-(3-carboxymethoxyphenyl)-2-(4-sulfophenyl)-2H-tetrazolium (MTS) to measure the ADC inhibitory concentrations (IC_50_), as DEspR inhibition alone is statistically insignificant in this assay. Previous data demonstrated that inhibition of DEspR enhanced MMAE efficacy in Panc1 and MiaPaCa2 cell lines (Fig. S6K,L); therefore, we identified MMAE as an appropriate payload. Since the ADC design includes release of free drug to measure bystander killing, we evaluated IC_50_ at 24, 48, and 72 hours. A progressive cytotoxic effect occurs across all PDAC cell lines (Panc1, MiaPaCa2, Capan-1, BxPC-3; Table 2, Fig. 3A), which likely reflects both initial targeting of the DEspR-positive cells, followed by drug release and targeting of the DEspR-negative cells by a bystander killing effect, as well as delays in cellular division.

**Fig. 3:**
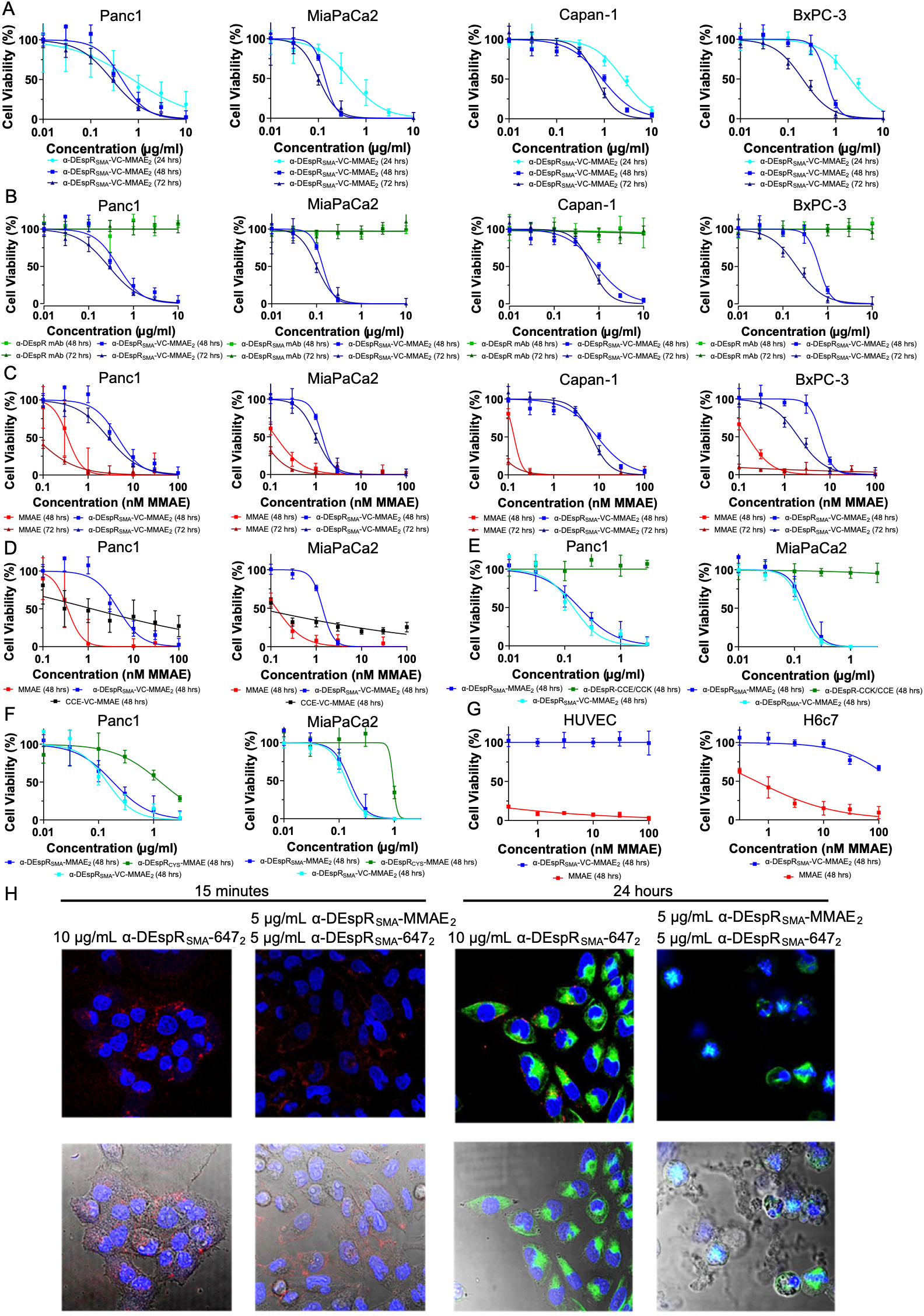
**ADC Cytotoxicity**. **(A)** Cytotoxicity of α-DEspR_SMA_–VC-MMAE_2_ in PDAC cells at 24hrs (light blue), 48hrs (blue), and 72hrs (dark blue). **(B)** Cytotoxicity of α-DEspR_SMA_–VC-MMAE_2_ [48hrs (blue), 72hrs (dark blue)] compared to α-DEspR mAb [48hrs (green), 72hrs (dark green)] in PDAC cells. **(C)** MMAE [48 hrs (red), 72hrs (dark red)], and α-DEspR_SMA_–VC-MMAE_2_ [48hrs (blue), 72hrs (dark blue)] cytotoxicity in PDAC cells, normalized to relative payload of MMAE. **(D)** Cytotoxicity of α-DEspR_SMA_–VC-MMAE_2_ (blue), CCE-VC-MMAE (black), and MMAE (red) at 48hrs in Panc1 and MiaPaCa2, with normalized MMAE concentrations. (**E)** Cytotoxicity at 48hrs of α-DEspR_SMA_–VC-MMAE_2,_ (light blue) α-DEspR_SMA_–MMAE_2_, (blue) and α-DEspR-CCK/CCE (green) in Panc1 and MiaPaC2 cells. **(F)** Cytotoxicity of α-DEspR_CYS_–MMAE (green), α-DEspR_SMA_–VC-MMAE_2_ (light blue), and α-DEspR_SMA_–MMAE_2_ (blue) at 48hrs in Panc1 and MiaPaCa2. Despite a higher DAR (3.68), α-DEspR_CYS_–MMAE was less effective in Panc1 (**IC_50,48h α-DEspRSMA-MMAE2_** 0.45 μg/ml, **IC_50,48hα-DEspRSMA-MMAE2_** 0.58 μg/ml, **IC_50,48hα-DEspRCYS-MMAE_** 4.26 μg/ml) and MiaPaCa-2 (**IC_50,48hα-DEspRSMA-MMAE2_** 0.14 μg/ml, **IC_50,48hα-DEspRSMA-MMAE2_** 0.16 μg/ml, **IC_50,48hα-DEspRCYS-MMAE_** 0.95 μg/ml). (**G)** Cytotoxicity at 48hrs of α-DEspR_SMA_–VC-MMAE_2_ (blue) and MMAE (red) in DEspR-negative HUVEC and DEspR-low H6c7 cells. **(H)** Representative confocal microscopy, comparing Panc1 cells treated with 10 µg/mL of 1:1 α-DEspR_SMA_–MMAE_2_ and α-DEspR_SMA_–647_2_ or 10 µg/mL α-DEspR_SMA_–647_2_ at 15min vs 24hrs. Cells were stained with NucBlue (blue, nucleus) and anti-tubulin-AF488 (green), allowing visualization of nuclear and microtubule structure.

**Table 2.** Cytotoxicity of ADC, Antibody, and Drug components in PDAC cell lines.

| Agent | IC<sub>50</sub>, 24 hr | IC<sub>50</sub>, 48 hr ([MMAE]) | IC<sub>50</sub>, 72 hr ([MMAE]) |
| --- | --- | --- | --- |
| **Panc1** |  |  |  |
| α-DEspR<sub>SMA</sub>-VC-MMAE<sub>2</sub> | 0.76 μg/mL (8.94 nM) | 0.45 μg/mL (5.29 nM) | 0.28 μg/mL (3.29 nM) |
| α-DEspR mAb | | >10 μg/mL | >10 μg/mL |
| MMAE |  | 0.25 ng/mL (0.35 nM) | 0.05 ng/mL (0.067 nM) |
| CCE-VC-MMAE |  | 6.68 ng/mL (1.21 nM) |  |
| α-DEspR<sub>SMA</sub>-MMAE<sub>2</sub> | | 0.58 μg/mL (6.9 nM) | |
| α-DEspR<sub>CYS</sub>-MMAE | | 4.26 μg/mL (50.4 nM) | |
| **MiaPaCa2** |  |  |  |
| α-DEspR<sub>SMA</sub>-VC-MMAE<sub>2</sub> | 0.45 μg/mL (5.29 nM) | 0.14 μg/mL (1.65 nM) | 0.10 μg/mL (1.18 nM) |
| α-DEspR mAb | | >10 μg/mL | >10 μg/mL |
| MMAE |  | 0.09 ng/mL (0.132 nM) | 0.04 ng/mL (0.062 nM) |
| CCE-VC-MMAE |  | 4.95 ng/mL (0.897 nM) |  |
| α-DEspR<sub>SMA</sub>-MMAE<sub>2</sub> | | 0.16 μg/mL (1.88 nM) | |
| α-DEspR<sub>CYS</sub>-MMAE | | 0.95 μg/mL (11.23 nM) | |
| **Capan-1** |  |  |  |
| α-DEspR<sub>SMA</sub>-VC-MMAE<sub>2</sub> | 2.51 μg/mL (29.53 nM) | 0.90 μg/mL (10.59 nM) | 0.68 μg/mL (8.04 nM) |
| α-DEspR mAb | | >10 μg/mL | >10 μg/mL |
| MMAE |  | 0.10 ng/mL (0.135 nM) | 0.03 ng/mL (0.041 nM) |
| **BxPC-3** |  |  |  |
| α-DEspR<sub>SMA</sub>-VC-MMAE<sub>2</sub> | 2.04 μg/mL (24.00 nM) | 0.64 μg/mL (7.53 nM) | 0.20 μg/mL (2.35 nM) |
| α-DEspR mAb | | >10 μg/mL | >10 μg/mL |
| MMAE |  | 0.11 ng/mL (0.156 nM) | 0.04 ng/mL (0.052 nM) |

Next, we compared the cytotoxicity of the α-DEspR_SMA_-VC-MMAE_2_ ADCs to α-DEspR mAb (Fig. 3B), and MMAE alone (Fig. 3C, normalized to payload of MMAE on the ADC). DEspR inhibition alone via the mAb is non cytotoxic at the tested dose range, allowing results from the MTS assay to reflect ADC function alone. MMAE rapidly kills all cells, in the low nanomolar to picomolar range, while cell death in PDAC cell lines requires a comparatively higher amount of ADC likely reflecting required ADC internalization by receptor positive cells, and then bystander killing effect. (Table 2)

Since SMA bioconjugation relies on non-covalent linkage, we next compared the cytotoxicity of the α-DEspR_SMA_-VC-MMAE_2_ to free CCE-VC-MMAE peptide. We anticipated that free CCE-VC-MMAE will possess a lower IC_50_ as cell entry is mediated by non-specific endocytosis rather than receptor-mediated binding. Free CCE-VC-MMAE is more potent than α-DEspR_SMA_-VC-MMAE_2_ at lower concentrations with lower IC_50_ in Panc1 and MiaPaCa2 cells. However, at 100 nM MMAE equivalence, cell viability is 27.2±14.1% for Panc1 and 25.5±6.2% for MiaPaCa2 for CCE-VC-MMAE treated cells, while it is 2.7±8.1% in Panc1 and 0.2±7.1% in MiaPaCa2 for α-DEspR_SMA_-VC-MMAE_2_ treated cells (Fig. 3D, Table 3). These results support α-DEspR_SMA_-VC-MMAE_2_ delivery following ADC internalization and free drug diffusion rather than pre-mature decoupling of CCE-VC-MMAE.

Given concerns about premature release with the valine-citrulline spacer in murine models,(*29*) we anticipated use of a glutamic acid-valine-citrulline spacer *in vivo* (α-DEspR_SMA_-MMAE_2_). Cytotoxicity is equivalent between α-DEspR_SMA_-VC-MMAE_2_ and α-DEspR_SMA_-MMAE_2_ in Panc1 and MiaPaCa2 cell lines (Figure 3E, Table 2). We also compared the α-DEspR_SMA_-VC-MMAE_2_ and α-DEspR_SMA_-MMAE_2_ ADCs to our commercial equivalent ADC, α-DEspR_CYS_-MMAE, to compare the effect of conjugation method. While α-DEspR_CYS_-MMAE has a higher DAR of 3.68, there is greater variability in drug loading and hinge reduction and conjugation with DMSO was required. α-DEspR_CYS_-MMAE possesses a higher IC_50_ in Panc1 and MiaPaCa-2 cells at 48 hours compared to either α-DEspR_SMA_-VC-MMAE_2_ or α-DEspR_SMA_-MMAE_2_ (Fig. 3F, Table 2).

To support the hypothesis that receptor-mediated internalization drives ADC toxicity, we evaluated cytotoxicity in DEspR-negative HUVEC and DEspR-low H6c7 cell lines. No significant toxicity is present in the DEspR-negative HUVEC cell line, and low toxicity (IC_50, 48h ADC_ >100 nM MMAE equivalence vs IC_50, 48h MMAE_ 0.65 nM) in the DEspR-low H6c7 cell lines, supporting receptor specificity of the ADC (Fig. 3G). Results from cell microscopy further support our hypothesis that receptor mediated internalization, followed by payload release, is the mechanism of ADC cytotoxicity. Upon treating Panc1 cells with a mix of α-DEspR_SMA_-VC-MMAE_2_ and α-DEspR_SMA_-647_2_ in a 1:1 ratio (total concentration 10 μg/ml), we tracked internalization over time and observed apoptosis from microtubule destabilization, the mechanism of cell death from MMAE, but not DEspR inhibition. Cells treated with 5 μg/ml of α-DEspR_SMA_-VC-MMAE_2_ and 5 μg/ml α-DEspR_SMA_-647_2_ undergo apoptosis or microtubule arrest in the majority of DEspR positive cells by 24 hours, while cells treated with only 10 μg/ml α-DEspR_SMA_-647_2_ show normal microtubule assembly (Fig. 3H). The presence of apoptosis or microtubule arrest in the majority of DEspR positive cells, but not DEspR negative cells (measured by signal of α-DEspR_SMA_-647_2_), supports the difference in observed IC_50_ at 24, 48, and 72 hours.

### SMA bioconjugate α-DEspR_SMA_-MMAE_2_ is specific for PDAC tumors

We next characterized the SMA conjugate α-DEspR_SMA_-MMAE_2_ *in vivo*, using an established model of orthotopic pancreatic peritoneal carcinomatosis.(*14,30*) To characterize ADC circulation, we injected tumor bearing rats 4 weeks after Panc-1 CSC tumor engraftment with either 0.1 mg/kg or 0.3 mg/kg intravenously, and collected rat serum for measurement of total ADC, total antibody, free peptide and free MMAE drug by ELISA and LC-MS. (Fig. 4A) The pharmacokinetic profile for α-DEspR_SMA_-MMAE_2_ fits a two-compartment model, with a rapid distribution half-life of 0.26 hours at 0.1 mg/kg and 0.32 hours at 0.3 mg/kg and an elimination half-life of 5.96 days at 0.1 mg/kg and 6.03 days at 0.3 mg/kg. The calculated maximum concentration, C_max,0.1mg/kg_ is 1488±71 and C_max,0.3mg/kg_ is 3890±484 ng/mL, which approximated the theoretical for each dose. The calculated AUC_0.1mg/kg_ is 97.52 µg/ml*h and the calculated AUC_0.3mg/kg_ is 324.14 µg/ml*h. Circulating non-conjugated antibody is not detectable until 24 hours after injection and remains low relative to the total amount of α-DEspR_SMA_-MMAE_2_, with levels higher in 0.3 mg/kg vs 0.1 mg/kg treated rats. Only at 72 hours after drug administration is minimal free MMAE present, while free CCE-MMAE is not detected at the limits of detection by RP-HPLC (Fig. 4B,C). In contrast, at 0.1 mg/kg dosing of α-DEspR_CYS_-MMAE, the C_max_ (1593±67 ng/ml vs. 1488±71 ng/ml) and distribution half-life (0.16 vs 0.26 hours) are similar, but the elimination half-life is substantially shorter (1.39 vs. 5.96 days), and the AUC is lower (10.84 µg/ml*h vs 97.52 µg/ml*h) than the SMA conjugate (Fig. 4D). Rapid clearance of α-DEspR_CYS_-MMAE is observed for both the total ADC as well as total mAb, reflecting accelerated elimination of α-DEspR_CYS_-MMAE rather than linker cleavage or premature MMAE release. Furthermore, α-DEspR_CYS_-MMAE levels fall below assay detection by 7 days.

**Fig. 4.**
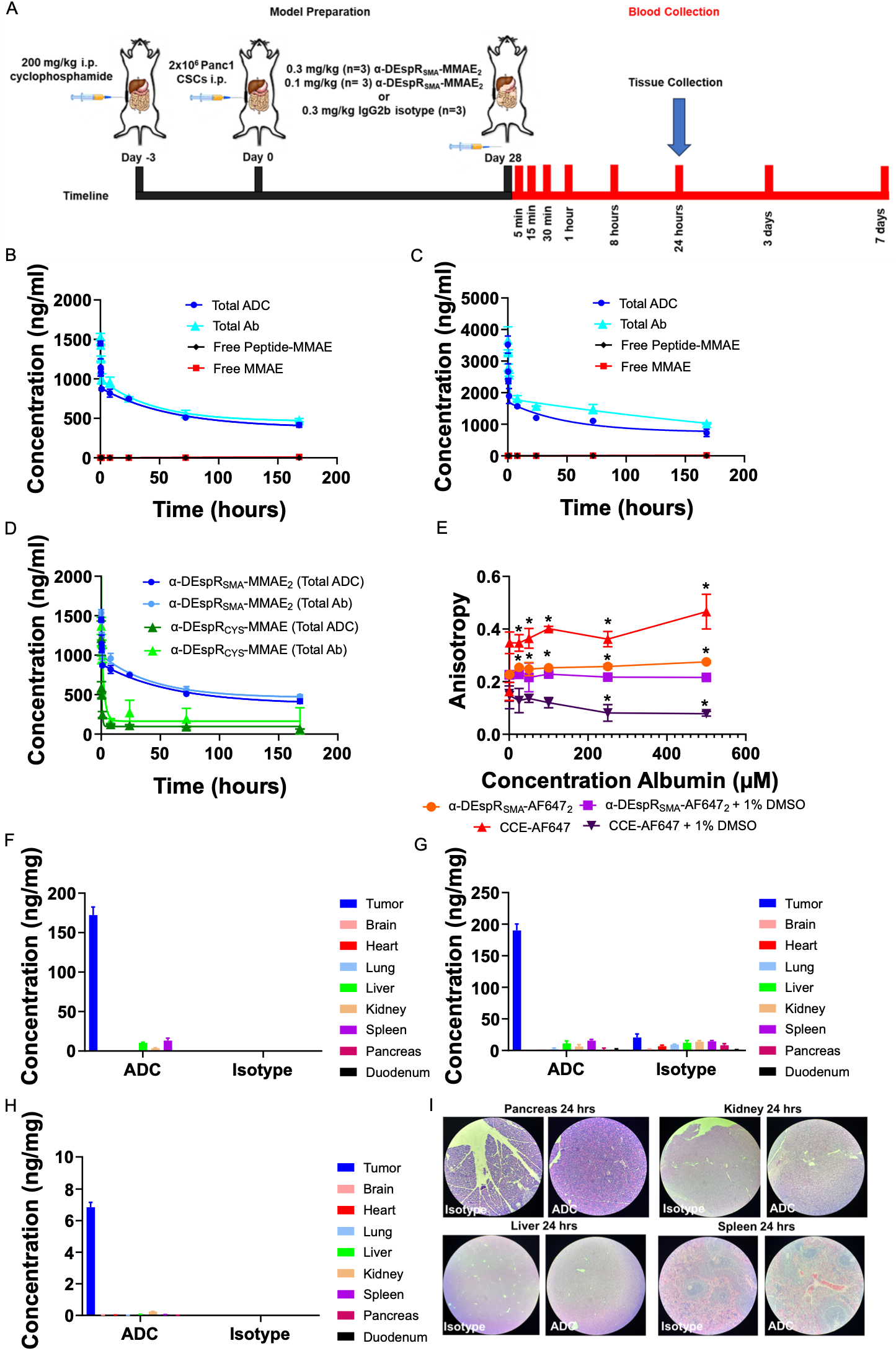
ADC Pharmacokinetics and Biodistribution Data. **A)** Representation of pharmacokinetic and biodistribution model (n=3/treatment group). **B)** Pharmacokinetics of 0.3 mg/kg IV of α-DEspR_SMA_–MMAE_2_, with a C_max_=3894-4820 ng/mL, t_1/2α_=0.32hr, and t_1/2β_=144.6 hrs. **C)** Pharmacokinetics of 0.1 mg/kg IV of α-DEspR_SMA_–MMAE_2_, with a C_max_ =1280-1490 ng/mL, t_1/2α_=0.26hr, and t_1/2β_=143.1hrs. No significant free MMAE or MMAE-bound peptide was detected for either group. **D)** A comparison of the pharmacokinetics of 0.1 mg/kg IV of α-DEspR_CYS_–MMAE and α-DEspR_SMA_–MMAE_2_. α-DEspR_CYS_–MMAE demonstrates a comparable C_max_ (1593±67 ng/ml vs. 1488±71 ng/ml) and t_1/2α_ (0.16hrs vs 0.26 hrs), but a shorter t_1/2β_ (1.39 days vs. 5.96 days) and lower AUC (10.84 µg/ml*h vs 97.52 µg/ml*h). **E)** Fluoresence anisotropy evaluating the interaction of CCE-AF647 and α-DEspR_SMA_–647_2_ with albumin. Addition of 0.1% DMSO, which corrects for AF647-albumin binding, shows that free CCE-AF647 weakly interacts with albumin (dark purple, K_d_=185 mM), but does not dissociate from α-DEspR_SMA_–647_2_ (purple). (**F-G)** Biodistribution of α-DEspR_SMA_– MMAE_2_ **(F)** total ADC, **(G)** total antibody levels, and **(H)** free MMAE concentration. **I)** H&E tissue sections where ADC, antibody, or MMAE signal was >1%. There was no overt histologic evidence of drug-related injury and no difference in morphology between ADC and isotype treated tissue.

Since we do not observe detectable levels of peptide dissociation in either *in vitro* or *in vivo* experiments, we evaluated whether dissociated peptide is bound to albumin. Using fluorescence anisotropy, we first evaluated the interaction of CCE-AF647 with albumin. Initially, the baseline anisotropy of 0.16±0.03 increases to an anisotropy of 0.47±0.06 with 500 µM albumin indicating tight restriction of the fluorophore, as this exceeds the theoretical anisotropy given by the Perrin-Jablonski equation for one-photon excitation (Fig. 4E). We hypothesized that this arises via binding of the AF647 dye in the albumin pocket rather a strong affinity between CCE-AF657 and albumin, as this pattern suggested dye immobilization.(*31*) To test this hypothesis, we added 1% DMSO to the solutions, which disrupts AF647 trapped within dye pockets. The anisotropy decreases to 0.078±0.009 at 500 µM albumin, suggesting a weak K_d_ of 185 mM albumin based on the modified Langmuir’s isotherm equation and greater rotational freedom with this CCE-albumin construct (Fig. 4E). Since these albumin concentrations are within the range of serum concentration, and dissociation could be observed even with this weak association, we next evaluated the anisotropy of α-DEspR_SMA_-647_2_ in albumin after 24 hours of equilibration. The anisotropy slightly increases from 0.225±0.003 for free α-DEspR_SMA_-647_2_ to 0.275±0.005 when mixed with 500 µM albumin (Fig. 4E). Again, we investigated whether the dye is being sequestered within albumin binding pockets and influencing the result. The values for the anisotropic difference disappear with addition of 1% DMSO (Fig. 4E). These data are consistent with the results from denaturing RP-HPLC, which do not detect free-MMAE separate from albumin.

We next assessed the biodistribution of α-DEspR_SMA_-MMAE_2_ twenty-four hours after intravenous injection in tumor bearing animals. This time point occurs after distribution equilibration, allowing accurate reflection of ADC distribution. We collected tissue from the brain, heart, lung, liver, kidney, pancreas, spleen, duodenum, and tumor, and quantified antibody and drug concentration via ELISA and LC-MS analysis, respectively. Both total ADC and total antibody concentrations are highest in the tumor (172.2±10.2 ng and 190.1±10.3 ng normalized to total protein), with low levels detected in the liver, kidney, and spleen (Fig. 4 F,G). Total ADC and total antibody concentrations of α-DEspR_SMA_-MMAE_2_ treated rats are similar to relative isotype levels in these tissues. Whereas isotype uptake occurs in the brain, heart, lung, pancreas, and duodenum, α-DEspR_SMA_-MMAE_2_ is not detected in these organs. Free drug is present predominantly in the tumor tissue (5.98±0.85 ng normalized to total protein), with minimal detection in the liver, kidney, and spleen, (0.08±0.01 ng, 0.25±0.01 ng, and 0.06±0.01 ng normalized to total protein), and not detected at the threshold of LC-MS in other organs. (Fig. 4H).

While levels of ADC and free drug in the liver, spleen, kidney, and pancreas are low, indicative of a high safety margin, we performed histology to assess potential toxicity from the ADC in healthy tissue. Hematoxylin and eosin (H&E) stain analysis of these organs shows no morphologic changes suggestive of cellular injury (Fig. 4I, Fig. S7). We confirmed organ integrity by evaluating for free caspase-3 as a surrogate for apoptosis and the results are comparable to the isotype control in these organs (Fig. S8).

### SMA bioconjugation is safe and non-immunogenic in immunocompetent rats

To further determine the safety and tolerability of α-DEspR_SMA_-MMAE_2_, we administered 10 times the anticipated treatment dose, 1 mg/kg, in immunocompetent RNU heterozygous rats. We injected 1 mg/kg intravenously weekly for 4 weeks, and monitored the animals for an additional 4 weeks. No clinical toxicity is noted in treated rats compared to saline controls, and hematologic data show no change in leukocyte, neutrophil, lymphocyte, red blood cell, or platelets circulating platelet number. (Fig. 5A-E). Hematoxylin and eosin stains of treated rat tissues from the biodistribution and safety studies to the saline and isotype controls reveal no overt organ injury after 8 weeks of treatment between ADC and saline controls (Fig. 5F). DAB staining of caspase-3 confirms a lack of treatment-induced cellular apoptosis in harvested organ tissue (Fig. 5F)

**Fig. 5.**
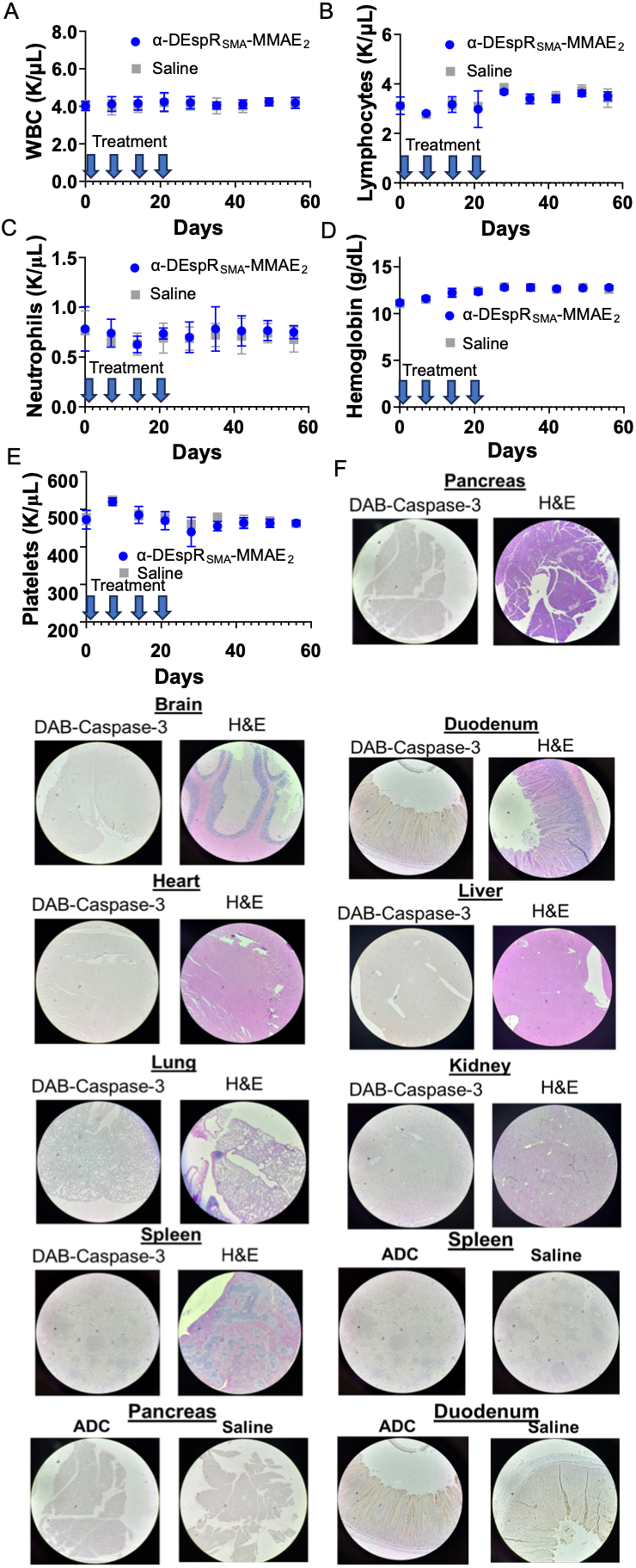
**ADC Safety Data**. Hematologic toxicities were screened in immune competent rats using 10x the anticipated treatment dose of α-DEspR_SMA_-MMAE_2_ (1 mg/kg, n=3 per group). No impact was noted in **A)** leukocytes, **B)** lymphocytes, **C)** neutrophils, **D)** Hemoglobin, or **E)** platelets after 4 treatment doses and following 4 weeks after treatment (α-DEspR_SMA_-MMAE_2_ = blue, saline = gray). **F)** Tissue was collected from saline and α-DEspR_SMA_-MMAE_2_ treated rats; in α-DEspR_SMA_-MMAE_2_ treated rats, no overt histologic injury or increased caspase-3 was noted in brain, heart, lung, liver, or kidney, and only minimal caspase-3 levels were noted in the duodenum (near the luminal surface), spleen (near the germinal centers), and near the borders of the pancreas, which were similar to saline treated rats.

Given the possibility of adverse acute inflammatory responses to the coiled-coil motif, we preliminarily assessed the innate and humoral stimulation to α-DEspR_SMA_-MMAE_2_. We administered 1 mg/kg α-DEspR_SMA_-MMAE_2_ intravenously weekly for 4 weeks in immunocompetent RNU heterozygous rats, and observed the animals for 4 weeks. ELISA measurements of serum IL-6 and TNF-alpha levels show no significant increase above vector control (Fig. 6A-B). We next evaluated humoral immune response to α-DEspR_SMA_-MMAE_2_, in order to evaluate the effect of the peptide coils on antigen presentation and antibody production. We compared anti-drug antibodies between native antibody and α-DEspR_SMA_-MMAE_2_. Detectable IgM levels are present starting at 6 weeks, while significant IgG levels are detected from 7 weeks onward. There is no statistically significant difference between anti-drug antibodies levels of native antibody and α-DEspR_SMA_-MMAE_2_ (Fig. 6 C,D).

**Figure 6.**
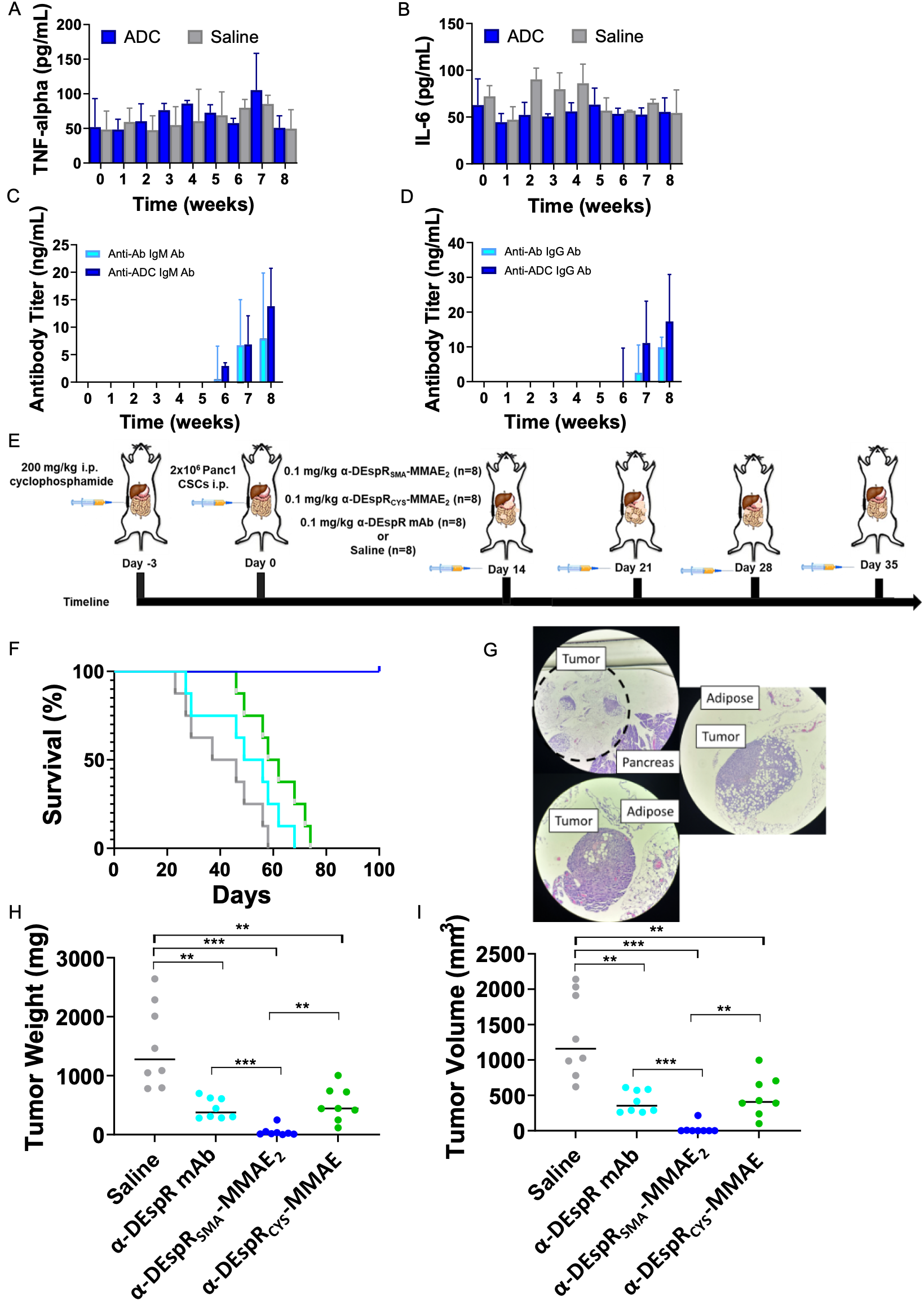
**ADC Immunogenicity and Efficacy**. **(A-B)** Comparison of **(A)** TNF-alpha and **(B)** IL-6 levels in heterozygous RNU rats treated with 1 mg/kg of α-DEspR_SMA_-MMAE_2_ (blue, n=3) and saline (gray n=3) during 4 weeks of treatment and 4 weeks of subsequent monitoring. **(C,D)** Comparison of **(C)** IgM and **(D)** IgG anti-drug antibodies from α-DEspR mAb (light blue, n=3) or α-DEspR_SMA_-MMAE_2_ (blue, n=3) during 4 weeks of treatment and 4 weeks of subsequent monitoring. **E)** Schema of pancreatic peritoneal carcinomatosis model. **(F)** Survival curve of PDAC treated rats. Median survival of therapies were: saline: 41.5 days, α-DEspR mAb: 52.5 days, α-DEspR_CYS_-MMAE: 60 days, α-DEspR_SMA_-MMAE_2_: 100 days. Mantel-Cox test demonstrated significant difference between all groups (p<0.0001), and Wilcoxon test showed a significant difference between α-DEspR_SMA_-MMAE_2_ and saline (p=0.0002), α-DEspR mAb (p=0.0002), and α-DEspR_CYS_-MMAE (p=0.0002). **G)** Histologic sections of grossly “tumor negative” α-DEspR_SMA_-MMAE_2_ treated rats, demonstrating residual tumor within the peri-pancreatic mesentery with a sparse extracellular matrix and minimal intratumoral capillaries. (**H,I)** Comparison of **(H)** total tumor weight and **(I)** tumor weight at time of euthanasia. Saline treated rats had higher tumor weights (1513±714 mg) and volume (1349±598 mm^3^) compared to α-DEspR mAb (444±174 mg, 410±156 mm^3^), α-DEspR_CYS_-MMAE (516±289 mg, 488±285 mm^3^), and α-DEspR_SMA_-MMAE_2_ (49±82 mg, 28.2±76 mm^3^).

### Anti-DEspR-(SMA-MMAE)_2_ ADC enhances survival in the peritoneal carcinomatosis model

Finally, we compared the efficacy of α-DEspR_SMA_-MMAE_2_ to the native antibody (α-DEspR mAb), and our commercially equivalent ADC using cysteine-maleimide coupling (α-DEspR_CYS_-MMAE), using an established model of pancreatic peritoneal carcinomatosis (Fig. 6E). Intraperitoneal treatment with anti-DEspR antibody, α-DEspR mAb, enhances survival at a dose of 1 mg/kg twice weekly(*14*); therefore, we selected a treatment dose of 0.1 mg/kg intravenously for all treatment arms to establish whether this conjugation method enhances survival. Based on our pharmacokinetic and pharmacodynamic data, we anticipated sufficient total drug exposure with weekly dosing at this treatment dose for four weeks to see an effect at 0.1 mg/kg with 8 rats. Increased overall survival is a primary outcome, while reductions in tumor mass and tumor volume are secondary outcomes.

Treatment with α-DEspR_SMA_-MMAE_2_ affords 100% survival at 100 days, while median survival with α-DEspR mAb is 52.5 days, and 60 days for α-DEspR_CYS_-MMAE. (Fig. 6F). Treatment with α-DEspR_SMA_-MMAE_2_ improves survival compared to α-DEspR_CYS_-MMAE (p=0.0002), α-DEspR mAb (p=0.0002), and saline (p=0.002). Treatment α-DEspR_CYS_-MMAE improves survival compared to saline (p=0.008), but not to α-DEspR mAb treatment. Treatment with α-DEspR_SMA_-MMAE_2_ eliminates visible tumor burden in 3/8 rats, though histologic sectioning of the tumors shows mesenteric invasion and small peri-pancreatic tumors in sectioned “normal” peripancreatic tissue (Fig. 6G). We euthanized rats for weight loss and poor body score (18/24 rats) or unresolving jaundice (6/24 rats), which correlated with either tumor invasion into the stomach or proximal duodenum causing gastric outlet obstruction (12/18 rats), invasion into the intestines and vasa recta (6/18 rats) or tumor invasion into the porta hepatis or liver (6/6 rats), with one rat noted to have evidence of early perforation at time of euthanasia (Fig. S9 A-H).

Treatment with α-DEspR_SMA_-MMAE_2_ decreases tumor weight at time of euthanasia, compared to saline (p=0.0006), α-DEspR mAb (p=0.0002), and α-DEspR_CYS_-MMAE (p<0.0022), with evidence of smaller tumors localized either near the pancreas or adjacent mesentery (3/8 rats), within the intestinal mesentery without invasion into the parenchyma (4/8 rats), or invading into the left lobe of the liver (1/8 rats) (Fig. 6H). Tumor weight of α-DEspR_CYS_-MMAE treated rats is comparable to α-DEspR mAb at time of euthanasia, without a significant difference in overall survival. Tumor volume also decreases at time of euthanasia in α-DEspR_SMA_-MMAE_2_ treated rats, compared to saline (p=0.0004), α-DEspR mAb (p<0.0001), α-DEspR_CYS_-MMAE (p<0.0023). Individual tumor volume in α-DEspR_SMA_-MMAE_2_ treated rats are smaller, reflecting both decreased individual tumor size and abundance of tumors.

## DISCUSSION

ADCs are a promising drug class, offering both a means of combination therapy and targeted drug delivery. An effective ADC design requires a tumor specific target, effective conjugation methods that preserve antibody binding and pharmacokinetic profiles, complementary drug selection, and appropriate linker design. Conjugation plays a pivotal role in this design consideration, as ill-defined antibody modification undermines ADC specificity and efficacy. Here, we show the importance of conjugation design in tandem with target selection to generate efficacious ADCs, while supporting a new method using heterotetrameric coiled-coil structures for site-specific conjugation.

DEspR is an ideal target for PDAC ADC design: human PDAC tumors exhibit high DEspR expression, while expression on normal tissue is low, and anti-DEspR therapy is efficacious in models of pancreatic peritoneal carcinomatosis, albeit at a higher dose. Further, DEspR inhibition decreases tumor vasculoangiogenesis and collagen production, preventing further tumor growth and reducing tumor desmoplasia. At the cellular level, DEspR also impacts tumor cell anoikis resistance and stress-survival, and reduced Mcl-1 protein expression, which enhances the effect of microtubule inhibition and destabilization.(*32,33*) This latter DEspR mechanism of action is relevant herein, and, thus, we selected a potent microtubule inhibitor (MMAE) for conjugation to DEspR. MMAE demonstrates a synergistic effect with DEspR inhibition with a low IC_50_ in PDAC cells.

The conjugation method plays a critical role in ADC efficacy. Use of conventional cysteine-maleimide conjugation results in an ADC, α-DEspR_CYS_-MMAE, which does not statistically improve survival nor reduce tumor burden compared to the native antibody in our *in vivo* rat model of pancreatic peritoneal carcinomatosis (Fig. 6F-I). Switching to a method of site-specific modification affords a drastic improvement in ADC efficacy, despite a lower DAR (2 vs 3.7), with long-term survival to 100 days. We attribute this performance difference to the more favorable pharmacokinetic profile of the heterotetrameric coiled-coil based site-specific modification, as the half-life of α-DEspR_SMA_-MMAE_2_ being 5-6 days ensures a greater drug exposure between doses and a higher steady-state dosing than α-DEspR_CYS_-MMAE, with a half-life of 1-2 days.

This finding is consistent with previous studies, which demonstrated the limitations of non-specific conjugation.(*34,35*) ADCs produced from cysteine hinge conjugation exhibit more rapid clearance compared to site-specific conjugation,(*36*) because the product mixture contains high DAR species, which are more prone to aggregation as well as being scavenging by reticuloendothelial system.(*5,6*) The resulting accelerated elimination lowers overall drug levels and increases the percentage of circulating ADCs with a lower DAR: both of these consequences lead to subtherapeutic drug levels.(*5,6,34,36*) Practically, this translates to inefficacious ADC therapy. The rapid decline in both total ADC and total antibody of α-DEspR_CYS_-MMAE supports this over premature linker or drug release alone, as in the latter case, total antibody levels would be higher and more closely approximate native antibody or our SMA conjugate. Additionally, cysteine hinge reduction negatively impacts IgG2b stability (*37*) which manifests as decreased *in vitro* efficacy after 48 hours in cell culture compared to the heterotetrameric coiled coil-based ADCs. Collectively, these results highlight the importance of the conjugation methods when selecting new ADC targets, as use of a non-specific conjugation shows no benefit over the native antibody.

We selected a supramolecular assembly approach over traditional methods of site-specific conjugation for several reasons. First, this method achieves greater product uniformity as there is only one conjugation site for the drug-loaded docking peptide to the antibody at the C-terminal position containing the receiving peptides. The tetrameric coil possesses a low K_d_ and preferentially forms in the presence of equimolar CCE and CCK with no production of dimer, trimer, or other such structures. This result contrasts with current methods of lysine, cysteine hinge, enzymatic, peptide affinity, or modified amino acid conjugation methods, which may have high affinity per conjugation site, but will naturally produce product variability equal to the number of possible conjugation sites.(*38*) Second, heterotetrameric coiled-coil conjugation occurs under mild, aqueous conditions, without requiring complicated conjugation clean-up, significant buffer exchanges, or the use of organic-aqueous solutions. Immunoglobulins are highly susceptible to thermal and chemical denaturing, and the addition of organic solvents or temperature modifications during conjugation can negatively impact ADC stability, target affinity, pharmacokinetics, and shelf-life.(*39*) Similarly, complex or multi clean-up steps result in significant yield reduction of the ADC.(*40*) As the antibody is the most expensive and delicate component in ADC design, minimizing reactions and handling is advantageous. Finally, the design must be compatible with different drug conjugation chemistries and flexible for future ADC compositions. Since drug conjugation occurs only on the CCE peptide, the composition of *f’* residue changes based on conjugation needs. This capability provides access to a “plug-and play” method of ADC design via assembly of a CCK-antibody with different drug-CCE conjugates where different drugs or linker chemistry are present in the conjugate. This approach also avoids the typical limitations of unnatural amino acid conjugation, as we do not require a prokaryotic vector for antibody production. Overall, this strategy provides a complementary method of site-specific conjugation with minimal impact to the native antibody along with diverse customization.

While supramolecular coiled coils are finding increasing utility in biomaterials and drug delivery platforms,(*18–20*) as well as for engineering a tumor activation switch and logic into antibodies(*41,42*), and recently antibody drug modifications,(*43–46*) this study offers several key advancements. First, this is the first application of a heterotetrameric coiled coil design. We selected the heterotetrameric linkage, and optimized the structure as it offers several key improvements over other oligomeric states. Based on the Crick Model of coiled coil assembly, higher oligomeric states increase the number of van-der-Waals interactions in the hydrophobic core, enhancing strength and stability, allowing a shorter sequence to achieve greater stability than lower oligomeric states.(*47,48*) This feature reduces the impact of conjugation on the antibody, allowing minimal negative impact on pharmacokinetic and pharmacodynamic properties. Furthermore, the use of a tetrameric design ensures an “all or nothing” conjugation design, as the presence of one pair of peptides on the C-terminus ensures that the conjugation products are either the ADC or the native antibody. The use of dimers or trimers can lead to a distribution of products, depending on composition, and a design limitation we sought to improve upon. The use of a higher order structure also increases “shielding” of the hydrophobic core, preventing acute inflammatory responses and likely explaining the lack of significant elevation of TNF-alpha and IL-6 levels.(*49*) Modifications of antibodies inherently affect the immunogenicity, safety, and longevity of ADC therapy, and remains an important concern with ADC design. While dimeric approaches have been explored(*43–45*), these specific issues were not addressed, nor did they demonstrate superiority to conventional conjugation strategies.

We do note several limitations to this study, which will need to be addressed prior to clinical translation. First, the selected antibody is a chimeric, murine IgG2b, which binds human, but not rodent DEspR. Therefore, generated ADCs should preferentially accumulate within the tumor, which expresses human DEspR, with minimal accumulation anticipated in macrophage rich organs, such as the liver and spleen, where most ADC clearance occurs.(*5,6*) This approach allows an easier direct comparison of pharmacokinetics between conjugation methods, but ignores off-target effects from receptor-mediated internalization by DEspR positive healthy cells. Since DEspR expression in non-PDAC tissue is low(*10,11*), we deemed this an acceptable approach to study SMA conjugation. Reassuringly, the SMA ADCs maintain high tumor specificity, with anticipated minimal accumulation in macrophage rich organs (liver and spleen), not exceeding isotype controls (a surrogate of Fc receptor binding). Low MMAE concentrations were detected in filtration organs (liver, kidney, and spleen), consistent with anticipated drug clearance.*(5,6* Therefore, SMA conjugation preserves binding affinity with high-intratumoral concentration and robust treatment outcomes. The use of murine IgG2b also decreases immunogenicity from the antibody, as repeat dosing of humanized anti-DEspR antibodies resulted in diminished outcomes overtime, likely related to neutralizing anti-drug antibodies. Future studies will require the examination of humanized anti-DEspR ADCs. Finally, the preclinical model for this study does have several limitations, such as more rapid tumor growth than human tumors, and smaller tumor volume than human disease. We selected this established model because it is reproducible, avoids the development of pancreatic insufficiency, which confounds weight loss, recapitulates a desmoplastic tumor environment, and generates tumor DEspR levels equivalent to human PDAC expression.(*11*) Furthermore, there is inherent variability with survival as an outcome, as tumors can lead to gastric outlet obstructions, perforations, or significant jaundice. Despite these limitations, treatment with the SMA ADC meaningful delays tumor growth and significantly improves survival outcomes.

Collectively, these data support both the exploration of anti-DEspR ADC therapy in PDAC and the further investigation of SMA strategies for drug development. PDAC survival outcomes critically lag behind those of other common tumors, owing to a lack of targeted, effective therapies.(*9,12*) Biologic therapy is grossly ineffective in PDAC(*9,12*) and there are currently no FDA approved ADCs for PDAC.(*9,49*) Effective treatments of PDAC require innovative approaches to address its unique biophysical and biochemical characteristics.(*50*) DEspR addresses both key PDAC survival and metastatic pathways, while also addressing tumor desmoplasia, a key consideration in enhancing biologic penetrance.(*14,15,50*) The impact on cellular stress pathways also synergizes well with microtubule inhibitors (e.g., MMAE), offering an avenue for synergistic ADC design. This non-covalent conjugation strategy, employing heterotetrameric coiled-coils, affords a plasma stable ADC of high product uniformity containing two MMAE drug molecules. The favorable pharmacokinetic profile and *in vivo* performance of the α-DEspR_SMA_-MMAE_2_ compared to the conventionally prepared ADC bodes well for its further development. SMA bioconjugation strategies are versatile, programmable, and compatible with complex biologics opening avenues to new therapies for cancer including pancreatic cancer – a disease in dire need of new treatments.

## Acknowledgements

This work was supported, in part, by NIH (T32 EB006359, M.W.G.; F30 CA220843 C.M.G.), Department of Gastroenterology and Hepatology at Yale University, the PhRMA Foundation Predoctoral Fellowship in Drug Delivery (A.R.), and the William Warren Professorship (M.W.G.). We also thank Drs. Herrera and Ruiz-Opazo at BU for discussions on DEspR, and Dr. Fred Gorelick at Yale for discussions on cell biology and pancreatic cancer.

## Author Contributions

C.M.G. and M.W.G. conceived the study. C.M.G. S.M.B., A.H. and A.R. prepared the antibodies and labeled coiled-coils. C.M.G. performed the *in vitro* assays and characterization. C.M.G executed the *in vivo* experiments. M.W.G. and W.M.S. provided laboratory space and resources. C.M.G. wrote the original draft of the manuscript. M.W.G. and C.M.G. secured funding. M.W.G. supervised the study. All authors contributed to the writing, figure preparation. and editing of the final manuscript.

## Supplementary Figures

**Fig. S1. (A-I).** Circular dichroism spectrum of all peptide acidic and basic pairings. Peptides are labeled by the hydrophobic amino acid in either the *e’* or *g* (first letter; L is leucine, V is valine, and I is isoleucine), and whether they are basic (second letter, K) or acidic (second letter E). **(J-L).** Circular dichroism of different acidic peptide pairings, demonstrating that mixtures of **(J)** L/E (red) and V/E (blue), **(K)** L/E (red) and I/E (blue), or **(L)** V/E (red) and I/E (blue) do not form supramolecular structures, likely because of charge repulsion between the species. **(M-O).** Circular dichroism of basic peptide pairings, demonstrating that mixtures of **(M)** L/K (red) and V/K (blue), **(N)** L/K (red) and I/K (blue), or **(O)** V/K (red) and I/K (blue) do not form supramolecular structures, as equimolar spectra (purple) are an average of the two species.

**Fig. S2. (A-C)** Analytical ultracentrifugation velocity scans, performed with 0.6 mg/mL of total peptide, using interference optics, were performed to analyze the molecular weight of CCE, CCK, and a CCE-CCK mixture, providing information of the formation of oligomeric states. **(A)** Velocity scans of peptide CCE, at demonstrating a single species with a low molecular weight. Using the modified Svedberg equation, this gives a molecular weight of approximately 4,000 kDa. **(B)** Velocity scans of peptide CCK, also demonstrating a single species with a low molecular weight. Using the modified Svedberg equation, this gives a molecular weight of approximately 4,000 kDa. **(C)** Velocity scans of a 1:1 mixture of peptide CCE-CCK, which again demonstrates a single predominant species; however, there is a shift in molecular weight. Using the modified Svedberg equation, this provides a calculated weight of 16,281 kDa (theoretical 15,788 kDa), supporting heterotetramer formation.

**Fig. S3. (A-C)** Analytical ultracentrifugation scans were performed using interference optics, with varying concentrations (0.2 mg/mL, left; 0.6 mg/mL, middle; 1.0 mg/mL, right) and varying rotational speeds (24,000 rpm, teal; 30,000 rpm, blue; and 54,000 rpm, purple), to provide more precise information on oligomeric species formation and strength of interaction between peptides. **(A)** Equilibrium scans of peptide CCK (top) with residuals (bottom), demonstrating a buoyant molecular weight (M_B_) of 4,288 kDa, with weak association (K_d_ = 0.0971 M), and fitting a single species model with good fidelity (RMSD of fit: 0.005281). **(B)** Equilibrium scans of peptide CCE (top) with residuals (bottom), demonstrating a buoyant molecular weight (M_B_) of 4,241 kDa, with weak association (K_d_ = 0.995 M), and fitting a single species model with good fidelity (RMSD of fit: 0.005442). **(C)** Equilibrium scans of equimolar peptide CCE-CCK (top) with residuals (bottom), demonstrating a buoyant molecular weight (M_B_) of 15,306 kDA, with strong association (K_d_ = 1.125 x 10^-10^), and fitting a tetrameric association model with good fidelity (RMSD of fit: 0.004801)

**Fig. S4. (A)** Thermal denaturing study of CCE-CCK, with representative spectrum shifts at 222 nM, ranging from 10°C to 90°C, which reflects the shift in the amide backbone and demonstrates the degree of retained coiled structure. At 90 ^°^C, only 18.2% ± 0.8% of the CCE-CCK construct unfolds. The repeat runs (blue, purple, and red), reflect the reversibility of this folding, demonstrating that aggregation does not form. **(B)** Chaotropic and thermal denaturing study of CCE-CCK, with representative spectra at 222 nm of samples equilibrated in 1M (light blue), 2M (light green), 4M (dark green), and 6M (black) guanidium chloride (GuCl), as temperature increases from 10°C to 90°C. Protein folding is observed, up to 53.5% at 90°C in 6M guanidinium chloride (GuCl). **(C)** Circular dichroism of peptides CCE-CCK in equimolar ratio at pH = 3, run with repeat thermal melting from 5°C to 90°C. While thermal melting was observed, this was reversible with cooling. **(D)** The impact of salinity was assessed by dialyzing the coil structure into different buffered solutions. As the salt concentration (measured as mM of sodium chloride) was decreased, the structure became less stable, ultimately collapsing without the presence of salt (dialysis of sample in deionized water).

**Fig. S5. (A)** SDS-PAGE (left non-reduced, right reduced), showing the difference in modified α-DEspR-CCK from α-DEspR antibody. **(B)** Representative chromatogram showing the modified CCE-vc-MMAE peptide after drug loading. **(C)** Representative chromatogram showing the modified CCE-EVC-MMAE peptide after drug loading. **(D)** Representative chromatogram showing the modified CCE-AF-488 peptide after drug loading. **(E)** Representative chromatogram showing the modified CCE-AF-647 peptide after drug loading. **(F)** LC-MS tracing confirming the molecular weight of the DBCO-EVC-MMAE linker. **(G)** ^13^C NMR spectra showing (top) the DBCO-EVC-MMAE linker (with glutamic acid protection), (middle) MMAE, and (bottom), the DBCO-EVC-PABC structure prior to addition of MMAE. **(H)** ^1^H NMR showing (top) the DBCO-EVC-MMAE linker (with glutamic acid protection) and (bottom) the DBCO-EVC-MMAE linker after deprotection. **(I)** Representative chromatogram showing the cysteine conjugated α-DEspR_CYS_-MMAE ADC, with peak for unconjugated mAb highlighted.

**Fig. S6. (A)** Representative UV-VIS spectrum of α-DEspR_SMA_647_2_ (DAR=2.00±0.14). **(B)** Representative UV-VIS spectrum α-DEspR_SMA_488_2_ (DAR=2.06±0.10). **(C)** Representative UV-VIS spectrum of α-DEspR_SMA_-VC-MMAE_2_ (DAR=2.08±0.15). **(D)** Representative UV-VIS spectrum of α-DEspR_SMA_-MMAE_2_ (DAR=2.10±0.13). **(E)** Representative UV-VIS spectrum of α-DEspR_CYS_MMAE (DAR=3.68±0.25). **(F)** Representative UV-VIS of α-DEspR mAb. **(G)** Representative RP-HPLC chromatogram of α-DEspR_SMA_-VC-MMAE_2_. **(H)** *Ex-vivo* stability assay of α-DEspR_SMA_-MMAE_2_, evaluating free MMAE in PBS (red) or rat serum (dark red) or CCE-MMAE release in PBS (blue) or rat serum (dark blue). No free drug and minimal free drug (<1%) were detected after 30 days. Free peptide was not detected in the samples using RP-HPLC. **(I,J)** Representative confocal microscopy of α-DEspR_SMA_647_2_ (red) internalization in **(I)** DEspR-low H6c7 and **(J)** DEspR-negative KV2 cells at 15 minutes vs 4 hours. Cells were stained with NucBlue (blue, nucleus) and Lysotracker (green, lysosome). Minimal α-DEspR_SMA_647_2_ uptake was noted in H6c7 cells and no uptake was noted in KV2 cells. **(K,L)** MMAE cytotoxicty at 24 hours were compared either with (red box) or without (hollow circle) 24hrs of α-DEspR mAb pre-treatment. Pre-treatment with α-DEspR mAb improved cytotoxicity of MMAE in **K)** Panc1 (IC_50,24h_: 0.365±0.04 vs 1.173±0.07 nM) and **L)** MiaPaCa2 cells (IC_50,24h_: 0.044±0.003 vs 0.260±0.020 nM).

**Fig. S7.** Representative H&E stains of rat organs for biodistribution study. **Fig. S8.** Representative Caspase-3 DAB staining of biodistribution organs **Fig. S9. Clinical Data of *In vivo* Efficacy Experiment. (A)** Clinical data of rats at time of euthanasia, with cause of death if euthanasia was required prior to 100 days. Common etiologies included gastric outlet obstruction (GOO), defined as visible evidence of tumor invasion into the proximal duodenum or antrum/pylorus of the stomach, perforation of gastrointestinal viscus, visualized with initial laparotomy, intestinal ischemia, defined as visible evidence of necrotic bowel with tumor invasion into adjacent vasculature, or persistent jaundice with evidence of invasion into biliary system. **(B-I)** Representative pictures of rats at time of euthanasia. (**B,C)** In α-DEspR_SMA_-MMAE_2_ treated rats with visible tumor, there were either **(B)** few solitary tumors **(C)** or small, hard to characterize tumors within the peripancreatic mesentery, confirmed with histologic staining. Tumor burden in **(D)** saline rats was comparable to **(E)** α-DEspR_CYS_-MMAE (tumor only) and **(F)** α-DEspR mAb treated rats. **(G)** Perforation with leakage of gastric contents noted at time of euthanasia. Salvaged bowel demonstrated a leak near a site of tumor invasion. **(H)** Tumor invasion into mesenteric vessels, leading to progressive weight loss and intestinal ischemia. **I)** Representative organs from 1/3 α-DEspR_SMA_-MMAE_2_ without visualized tumor at the time of euthanasia. Tumor was noted within the peri-pancreatic mesentery.

**Table S1. Comparison of current landscape of site-specific conjugation.**

Contemporary methods of site-specific conjugation are shown, with their ability to enable site-specific modification, uniform (100% conversion of ADC at all conjugation sites) drug loading, requirement of altered glycosylation of the antibody (glycosylation-dependent changes) to allow conjugation, and whether vector selection is not a factor in conjugation.

**Table S2.** Sequences of modified leucine-zipper sequences

Prepared sequences of each peptide, labeled for the *g*/*e’* site that was modified by a hydrophobic amino acid, followed by whether glutamic acid (E) or lysine (K) was used.

**Table S3.** Quality of Model Fit for Each Oligomeric State.

**Table S4.** Best predictive fit of each species and estimated dissociation constant

## MATERIALS AND METHODS

### Study Design

The aim of this study was to generate a site-specific method for ADC conjugation, which did not impact antibody glycosylation or vector selection, avoided the exposure of the antibody to organic solvents, and allowed conjugation with minimal clean up, using a bio-orthogonal method to available techniques. Using this technique, we hoped to optimize ADC development targeting a novel PDAC receptor, DEspR. Peptide design was performed *in silico*, using available data(*25*) and CC Builder 2.0, with optimization based on anticipated structures.(*24–26*) The most favorable peptide structures were prepared by Abclonal for *in vitro* testing. Following CD and AUC experiments, the CCK was recombinantly added to the C-terminus of an anti-DEspR antibody, and the CCE peptide was used for chemical conjugation. *In vitro* experiments were designed to characterize ADC cytotoxicity, safety, and estimate stability. *In vivo* experiments mirrored previously reported data on anti-DEspR monoclonal antibodies in PDAC. The ADC treatment concentration of 0.1 mg/kg was selected based on *in vitro* data comparing antibody and ADC efficacy and to better highlight differences between conjugation methods. Pharmacokinetic and biodistribution studies were performed prior to efficacy experiments to support adequate treatment doses, and safety and immunogenicity experiments were performed to characterize possible end-organ injury at 10x the anticipated treatment dose and possible increased immunogenicity from the conjugation method compared to the native antibody. Animals were randomized to treatment groups, based on weight at the time of the study, and no data was excluded from the analysis.

### Circular Dichroism Experiments

Peptides were prepared as 100 μM solutions and in Tris-buffered saline (TBS) pH 7.6, and samples were run as 5 replicates, using a 1 mm QS 100 (Hellma) cuvette. Samples were run on a CS/2 Chirascan (Applied Photophysics) CD spectrometer. Circular dichroism was measured in mDeg, and background corrected samples were converted to mean residue molar ellipticity (ϴ).

Helicity was determined from the following equation:

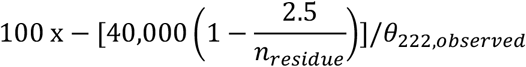

Thermal experiments were performed from 10°C to 90°C at a rate of 1°C/ min, with scans collected every 5°C. For repeat studies, the instrument was allowed to cool and equilibrate to 10°C, and a single CD run was performed to confirm no artifacts were present from heating. For denaturation studies, samples were equilibrated with varying concentrations of Guanidium Chloride (Sigma Aldrich). For pH studies, peptides were dissolved in pH adjusted TBS. For salt studies, peptides were dialyzed against DI water for 72 hours (6 dialysis volumes of 2 L to 4 mg total peptide). Samples were run at maximum concentration of tetramer (30 μM monomer), with increasing NaCl (Sigma, S7653) concentration.

### Analytical Ultracentrifugation

Sedimentation equilibrium (SV) and sedimentation velocity (SE) experiments were performed using six-centerpieces with interference optics in a Beckman XL-I. Interference optics were selected given the size and low aromatic content of each species. Samples for SV experiments were run at 40,000 rpm at 20°C using 0.6 mg/ml total peptide in TBS; SE experiments were run at 24,000, 30,000, and 45,000 rpm at 20°C using 0.2, 0.6, and 1.0 mg/ml total peptide in PBS. SV data was analyzed using Sedfit with a continuous c(s) distribution model.(*51*) A continuous sedimentation coefficient distribution model of the Lamm equation was used to analyze each species, with a maximum entropy model was introduced to reduce over-assumptions about the data.(*52–54*)

### Recombinant Antibody Preparation

Purified plasmid was mixed in a 1:2 ratio of heavy chain to light chain plasmids and transfected into Expi293 cells using the ExpiFectamine transfection kit. Cells were incubated for 7 days and then pelleted and the supernatant collected. The supernatant was filtered through a 0.22 µm filter, mixed in a 1:1 ratio with binding buffer (1.5 M Glycine/NaOH, 3 M NaCl, pH 9.0), and purified using rProtein A gravity columns. 0.2 M Glycine/HCl (pH 2.5) and 1 M Tris/HCl (pH 9.0) were used as elution and neutralization buffers, respectively. Antibodies were buffer-exchanged into PBS via spin filtering, aliquoted, and stored at-20 °C until further use.

### SDS Page

Purified antibodies were analyzed using Bio-Rad 4–15% precast gradient gels. For reduced samples the proteins were mixed with 4x Laemmli sample buffer containing 10% β-mercaptoethanol and heated at 95°C for 5 minutes. For non-reduced samples, β-mercaptoethanol and heating were omitted. Electrophoresis was performed at a constant voltage of 120V for approximately 60 minutes. Proteins were visualized by gel staining with InstantBlue protein stain solution.

### Chemical Synthesis

^1^H NMR and ^13^C NMR spectra were recorded on an Agilent 500 MHz VNMRS spectrometer in the stated solvents using tetramethylsilane as the internal standard. Chemical shifts were reported as parts per million (ppm) on the δ scale from the instrument standard. All materials were obtained from Sigma-Aldrich, and unless otherwise stated, were used without further purification. All solvents were purchased as ACS reagent grade or anhydrous if indicated. All reactions were carried out under a nitrogen atmosphere, unless otherwise indicated. DBCO-Val-Cit-PABC-MMAE and DBCO-Glu-Val-Cit-PABC-MMAE linkers were synthesized as previously reported by Anami et al.(*55*) CCE Peptide conjugation of AlexaFluor 488 DBCO (MedChemExpress), AlexaFluor 647 DBCO (MedChemExpress), synthesized DBCO-Val-Cit-PABC-MMAE or DBCO-Glu-Val-Cit-PABC-MMAE (2.0 eq) occurred in phosphate buffered solution (pH = 7.4) at room temperature for 4 hours while stirring. Following completion of the reaction, purification was performed with Amicon 3kDa MWCO spin column (Millipore Sigma), with three sequential washes at room temperature. The retentate was collected and analyzed via NanoDrop Ultra UV-VIS and RP-HPLC

### ADC Generation

Recombinant 7c5-CCK was mixed with labeled CCE peptides in a 1:2.1 ratio at 4°C for 4 hours. Following reaction, the ADC was purified by sequential washing in a Amicon 10kDa MWCO spin column. For conventional labeling studies, 7c5 or IgG2b isotype antibodies were labeled using Alexa Fluor 488 or 568 Antibody Labeling Kit (Fisher Scientific) following manufacturer instructions. Fluorophore loading was calculating using UV-VIS, with calculated loading of 2.2 and 2.4 respectively. Preparation of α-DEspR_CYS_-MMAE was performed based on previously established methods.(*28*)

### UV-VIS Experiments

0.1 mg/mL solutions of each ADC, were analyzed on a NanoDrop™ One Spectrophotometer UV-VIS, using PBS buffer (pH 7.4) for sample characterization and blanks. Extinction coefficients for the antibody were obtained based on a theoretical extinction at 280 (2.1 x 10^5^) and an experimentally calculated extinction at 248 (1.4 x 10^5^), based on Beer’s Law. DAR was calculated based on Beer’s Law, using reported extinction coefficients of MMAE or fluophores.(*56*)

### ADC HPLC Analysis

All Non-reductive RP-HPLC was performed using a Zorbak 300CN-SB column on a Shimadzu LC-2030C Plus HPLC, equipped with an RF-20A XS for fluorescence experiments and LC-MS experiments were performed using an Agilent 6546A QTOF3 with 1290 Infinity II LC operated in the positive ion model was used. Multiple-reaction monitoring (MRM) was performed to quantify MMAE and the innate standard. Loaded sample concentrations were 0.1 mg/mL for all ADC experiments. For all HPLC studies, three sample blanks were run prior to each sample and samples were run in triplicate and raw HPLC data was plotted into PRISM 11.

For ADC analysis, mobile phase A was 2.3 M ammonium sulphate (Sigma Aldrich) in 50 mM phosphate buffer (pH 6.8). Mobile phase B was 50 mM phosphate buffer with 30% methanol. The following protocol was used: 10% Mobile Phase B for 2 minutes, then increase to 80% Mobile Phase B for 10 minutes, hold for 5 minutes, then decrease to 10% Mobile phase B over 5 minutes, and hold for 5 minutes. For fluorophore-based experiments, the following protocol was used: 10% Mobile Phase B for 5 minutes, then increase to 60% Mobile Phase B for 10 minutes, hold for 5 minutes, then decrease to 10% Mobile phase B over 5 minutes, and hold for 5 minutes. For drug release studies, α-DEspR_SMA_-MMAE_2_ was prepared in aliquots of PBS or RNU serum. At 1 day, 3 days, 7 days, 14 days, 21 days, and 30 days, samples were prepared for free drug or CCE-Peptide analysis. For the former samples, protein precipitation was performed by addition of 3M NaCl (Sigma) followed by addition of cold methanol (4 equivalents to serum), incubated at-20°C for 2 hours. The sample was run with a Mobile Phase A of water with 0.1% trifluoroacetic acid (TFA) and a Mobile Phase B of acetonitrile with 0.1% TFA. Samples were run at 30% solvent B for 2 minutes, increased to 90% over 6 minutes, held for 2 minutes, then returned to 30% over 2 minutes, and held for 2 minutes. A standard curve was run from 1000 ng/mL to 0.5 ng/mL of MMAE.

For the latter, larger proteins were eliminated using a 30 kDA MWCO Amicon column, followed by ADC analysis RP-HPLC conditions. To evaluate for albumin-CCE aggregates, samples were run following use of a 100 kDA MWCO Amicon column. Here mobile phase A was 1% TFA in water. Mobile phase B was 1% TFA in acetonitrile. The sample was equilibrated with 30% Mobile Phase B for 5 minutes, increased to 60% Mobile Phase B for 20 minutes, held for 5 minutes, then decreased to 30% Mobile phase B over 10 minutes, and held for 5 minutes.

### Antigen Binding

For all ELISA assays, 96-well Nunc-Immuno MaxiSorp plates (Sigma Aldrich) were used. For antigen binding, plates were coated with 5 μg/ml of the antigen peptide (M_1_TMFKGSNE_9_) in bicarbonate buffer, and incubated overnight at 4°C. Plates were washed with PBS thrice, followed by blocking with 200 μl 1% bovine serum albumin (BSA) in PBS for 4 hours at 4°C. After washing, drugs were added in triplicates from 0.5 to 500 nM in 100 μl. After incubation for 1 hour at 37°C, the cells were washed thrice, and 50 μl of 1:2000 HRP-labeled anti-mouse IgG (Sigma) was added for 1 hour at 37°C. Cells were washed thrice, prior to addition of 100 μl of TMB substrate (BD) followed by stopped with 2M HCl. Optical absorbance was read at 450 nm.

### Cell Culture

Panc1, MIA PaCa2, Capan-1, BxPC-3, HUVEC, KV-2, BJ Fibroblasts, and mIMCD were purchased through ATCC and cultured per manufacturer instructions. H6c7 cells were purchased through Kerafast and cultured per manufacturer instructions. Cells were harvested at 60% confluence using trypsin-EDTA (0.25%) (Fischer Scientific); experiments were performed from passage 3-8.

### Flow Cytometry

Cells were labeled with 30 µg/ml α-DEspR_SMA_-488_2_, α-DEspR_LYS_-488, or IgG2b-488 antibody 20 minutes at 4°C. Detection was performed using an LSRII Flow Cytometer (BD Bioscience) with a 488 nm laser with 530/30 (505 LP) filter for AF-488.

### Anti-DEspR/MMAE Synergy Experiment

Panc1 or MiaPaCa2 cells were seeded at 2,000 cells per well in 200 ul media in 96-well clear bottom plates (Corning). After 48 hours, cells were either treated with 10 µg/ml of anti-DEspR antibody or media with equivalent PBS for 24 hours. Prior to treatment, plates were assessed by morphology with a Nexcelom Celigo Micro-Well Plate Imager before and after media exchange and drug loading to normalize for MMAE treatment effect. Cells were exposed to varying concentrations of MMAE (n = 5 per sample), and viability was assessed after 24 hours of treatment. Viability experiments were performing using a Nexcelom Celigo Micro-Well Plate Imager, with total cell count captured using NucBlue Live cell stain (ThermoFisher) and dead cells using Live/Dead Red stain (Invitrogen), using the instruments live/dead cell count strategy, with total cell count supported with brightfield images.

### Fixed Cell Fluorescent Microscopy

Immunofluorescence staining was performed as previously described.(*14*) Cells and were labeled with 10 µg/ml α-DEspR_SMA_-568_2_ for antibody internalization, or treated with 5 µg/ml of α-DEspR_SMA_-MMAE_2_ and for 5 µg/ml of α-DEspR_SMA_-568_2_ cytotoxicity experiments. For internalization experiments, cells were treated with lysotracker green (ThermoFisher). For microtubule inhibition, cells were fixed with 2% paraformaldehyde, permeabilized with Triton-X, and then stained with anti-tubulin-AF488 (ThermoFisher). For murine antibody internalization, imaging was performed with a Zeiss Axiostop fluorescence microscope, as previously described127 and confirmed with confocal studies using the Leica SP5, with 1 AU pinhole aperture, 0.7 µm Z-stacks taken for comparison. For antibody internalization, excitation was performed with 405 nm and 543 nm laser lines, emission was collected from 430-475 nm and 600-700 nm to prevent overlap. Quantification was performed using ImageJ with JACoP plugin-software.

### MTS Assay

Cells were seeded between 2,000-3,000 cells/ well in 200 ul media, depending on their doubling times. Once cells reached 20% confluence, varying concentration of the test reagent was added. Antibody and ADC samples were allowed to bind 15 minutes at 4°C, followed by removal of media, washing thrice with PBS, and then addition of pre-warmed media. After 24, 48h, and 72 hours, media was removed and 0.5 mmol MTT was added in serum free medium. After 3 hours, medium was removed and 50 ul of DMSO was added. Absorbance at 590 nm was measured. Cells did not exceed 80% confluence for assay, allowing direct measurement of cell viability before confluence was ever reached.

### Orthotopic Panc1 pancreatic peritoneal carcinomatosis model

Orthotopic pancreatic peritoneal carcinomatosis models were prepared as previously described.(*13,14*) Animal experiments were performed under IACUC protocols 2021-20428 and 2025-20428, which comply with government regulations (PHS and USDA) and the AAALAC. Selected endpoints differed from prior studies,(*14*) as death was deemed an inhumane endpoint. Acceptable endpoints as surrogates include weight loss of ≥ 10% from maximum, body score <2, persistent jaundice (lasting >24 hrs), persistent ascites (lasting >24 hrs), as well as additional health concerns as noted by veterinarian and husbandry staff. Animals were randomized to treatment group based on weight and palpable tumors at time of injection. Veterinarian and Husbandry staff were blinded to treatment groups when determining euthanasia.

### Pharmacokinetic Studies

For pharmacokinetic studies, tumor bearing rats were treated with 0.1 or 0.3 mg/kg of α-DEspR_SMA_-MMAE_2_ (n=3) after 4 weeks of tumor engraftment. Blood was collected at 5 minutes, 15 minutes, 30 minutes, 1 hour, 8 hours, 24 hours, 72 hours, 7 days, and 14 days after injection in 1% EDTA tubes. Serum was collected after centrifuging samples at 2,000x g for ten minutes, and collecting the supernatant.

### Biodistribution Studies

For pharmacokinetic studies, tumor bearing rats were treated with 0.3 mg/kg of α-DEspR_SMA_-MMAE_2_ or IgG2b isotype (ThermoFisher) (n=3) after 4 weeks of tumor engraftment. After 24 hours, rats were euthanized by terminal perfusion of the heart with saline. Tissue (brain, heart, lung, liver, pancreas, duodenum, spleen, kidney, and tumor) were collected. Tissue was either fixed in 10% buffered formalin for immunohistochemistry, or underwent protein extraction. For protein extraction, tissue was flash frozen, then digested by immersion of 50 mg of tissue in 500 μl of RIPA buffer and protease cocktail (ThermoFisher) with mechanical homogenization using Precellys 24 protein homogenization kit (Bertin Technologies) per manufacturer instructions. The supernatant was harvested, then re-centrifuged at 2000 xg for 15 minutes to remove any cellular debris. Tissue protein levels were assessed by absorbance at 280 nm with correction for nucleic acids (NanoDrop™ One Spectrophotometer).

### ADC Quantification

Total ADC and antibody were quantified using ELISA, based on modifications of previously established methods.(*57*) A murine anti-MMAE antibody (4 μg/mL) [BioRad] and synthesized antigen peptide (10 μg/ml of M_1_TMFKGSNE_9_) were used for analyte capture respectively, and detection was performed using HRP-conjugated goat anti-mouse IgG2b [Invitrogen] solution (100 ng/mL). Analysis was performed on a SpectraMax 7 Plate Reader.

### Fluorescence Anisotropy Experiments

Studies were performed on a SpectraMax 7 Plate Reader, using fluorescence and fluorescence anisotropy settings. Samples were run using black wall 96 well plates (Corning). Initial fluorescence readings were performed from titrations at 300 nM to 1 nM concentrations in 100 µl in triplicate. Anisotropy was calculated for samples over this range, with G calculated from the individual AF647 fluorophore. Background subtraction was performed for each sample. A concentration of 3 nM was selected for quantification of CCE-AF647 and α-DEspR_SMA_-647_2_ based on the appropriate signal to noise ratio. Samples were prepared in varying concentrations of albumin, and experiments were performed after 24 hours of mixing at 37°C. Samples contained either 1% DMSO or no DMSO to evaluate for interaction between hydrophobic binding pockets.

### Immunogenicity Experiments

Heterozygous RNU control rats (immunocompetent) were treated with 1 mg/kg of α-DEspR_SMA_-MMAE_2_, α-DEspR mAb, or saline (n=3) once a week for 4 weeks. Blood was collected prior to treatment and prior to each injection, as well as for a total of 4 weeks after treatment. Analysis of TNF-alpha and IL-6 levels were performed using a Legend Max Rat IL-6 ELISA (BioLegend) and ELISA MAX Deluxe Set Rat TNF-alpha (BioLegend) assay kit respectively. Total anti-drug IgM and IgG antibodies were detected using previously reported methods.(*58*) Plates were coated with either α-DEspR_SMA_-MMAE_2_ (4 μg/mL) or α-DEspR_SMA_ mAb and incubated overnight at 4 °C. Anti-IgG2b IgM rat (Novus Biologics) or anti-IgG2b IgG rat (Novus Biologics) were used as standards respectively. Detection was performed using either HRP-conjugated goat anti-rat IgM or HRP-conjugated goat anti-rat IgG (ThermoFisher). Signal read-out was collected using an absorbance signal at 450 nm on a SpectraMax 7 Plate Reader.

### Safety Experiments

Heterozygous RNU control rats were treated with 1 mg/kg of α-DEspR_SMA_-MMAE_2_, α-DEspR mAb, or saline (n=3) once a week for 4 weeks. Blood was collected prior to treatment and prior to each injection, as well as for a total of 4 weeks after treatment. Weekly blood collection was performed with a HEMAVET 950 FS Auto Blood Analyzer with rat-species settings. Tissue was collected at completion of the study for immunohistochemistry. Formalin fixed tissues were embedded and sectioned by the Yale Immunohistochemistry Core. H&E-stained slides were prepared of organs and analyzed for histologic evidence of cellular injury. DAB-Capase-3 slides were prepared for evaluation of drug-induced injury.

### Efficacy Study

Tumor bearing heterozygous RNU rats were randomized to 0.1 mg/kg of α-DEspR_SMA_-MMAE_2_, 0.1 mg/kg of α-DEspR mAb, 0.1 mg/kg of α-DEspR_CYS_-MMAE, or saline (n=8) after 2 weeks of tumor engraftment. Rats received weekly injections for 4 weeks, and were followed for survival surrogate endpoints as noted above. Tumor tissue was harvested at time of diagnosis and collected volume was obtained with calipers using a standard ellipsoid calculation. Sample weight was collected in aggregate as well, following tissue washing in PBS and drying to remove debris.

## Statistical Analysis

Statistical analyses were performed using GraphPad PRISM 11. Paired Student’s t-test was used to compare means between two groups. Chi-square tests-of-independence were used to compare categorical data. Analysis of variance and non-parametric Kruskal–Wallis one-way analysis of variance were performed when appropriate for ≥3 study groups. Correlation analysis was performed using the Pearson correlation coefficient (R) between continuous variables. Colocalization across imaging was performed using Manders coefficient. Differences in OS were calculated with the Kaplan-Meier survival curve, Mantel-Cox log rank statistic, and Holm-Sidak multiple comparison test. P values were corrected using the Bonferroni multiple comparison testing. Statistically significant values were indicated as follows: *p ≤ 0.05,**p ≤ 0.01,***p ≤ 0.001, and ****p ≤ 0.0001 unless otherwise stated.

