## Supplementary material for "Supramolecular Site-Specific Antibody Drug Conjugates Outperform Cysteine-Conjugated Analogs in Pancreatic Peritoneal Carcinomatosis": SI Text, Figures, and Tables

*Corresponding authors:

Christopher M. Gromisch

Room 401E

Mark W. Grinstaff

Room 519

590 Commonwealth Ave, Boston MA

Boston, MA 02215

**Analytical Ultracentrifugation:** Sedimentation equilibrium (SV) and sedimentation velocity (SE) experiments were performed using six-centerpieces with interference optics in a Beckman XL-I. Interference optics were selected given the size and low aromatic content of each species. Samples for SV experiments were run at 40,000 rpm at 20°C using 0.6 mg/ml total peptide in TBS; SE experiments were run at 24,000, 30,000, and 45,000 rpm at 20°C using 0.2, 0.6, and 1.0 mg/ml total peptide in PBS. SV data was analyzed using Sedfit[46] with a continuous c(s) distribution model. For SV experiments, buffer density was 1.0058 g/mL, and the vbar of each peptide was: CCE = 0.799 mL/g, CCK = 0.742 mL/g, and CCE-CCK = 0.762 mL/g, based on estimates from Sednterp. A continuous sedimentation coefficient distribution model of the Lamm equation was used to analyze each species:

$$\mathrm{Min}_{\mathrm{c(s)}}\ \{\sum_{i,j}[a(r_i,t_j) - \int c(s)\mathrm{L}(s,D(s),r_i,t_j)ds]^2\}$$

A maximum entropy model was introduced to reduce over-assumptions about the data[47]:

$$\mathrm{Min}_{\mathrm{c(s)}}\ \{\sum_{i,j}[a(r_i,t_j) - \int c(s)\mathrm{L}(s,D(s),r_i,t_j)ds]^2 + \alpha \int c(s)\ln c(s)\,ds\}$$

This model holds true under the following assumptions: either the species are non-interacting (which should fit for CCE and CCK) or if the other species are stable during sedimentation, indicating a higher $K_d$ and minimal dissociation (Ideal case for CCE-CCK).[48]

For calculation of the molecular weight of each sedimentation species, a modification of the Svedberg equation was used, by first calculating the friction coefficient, $f_o$, as defined as a a compact, ideal sphere of defined radius r, in the solvent of viscosity, η.

$$f_o = 6\pi\eta r$$

Substituted into the Svedberg equation, this gives:

$$\frac{M}{N_a} = \frac{sf}{1-vbar*p}\left(\frac{f}{f_o}\right)6\pi\eta\left(\frac{3}{4\pi}*\frac{M}{N_a}*\mathrm{vbar}\right)^{1/3}$$

This resolves to give the following equation:

$$\left(\frac{M}{N_a}\right)^{2/3} = \frac{s\left(\frac{f}{f_o}\right)}{1-vbar*p}6\pi\eta\left(\frac{3*vbar}{4\pi}\right)^{1/3}$$

This then provides the particle mass and partial specific volume of an ideal sphere, where M is the particle mass, $N_a$ is Avogadro's number, f is the fictional coefficient, vbar is the partial specific volume (which reflects variations in the volume from the composition of each particle), and $\rho$ is the density of the solvent. Diffusion was then calculated as follows.

$$D(s) = \frac{\sqrt{2}}{18\pi}kTs^{-1/2}(n(f/f_o)_w)^{-3/2}((1 - vbar*\rho)/(vbar)^{1/2}$$

SE data was analyzed using Sedphat[48,49] with single-species non-interacting fits, monomer-dimer, monomer-trimer, and monomer-tetramer fits, and heterotetramer fits. Temperature-corrected partial specific volumes (CCE= 0.7462 mL/g; CCK = 0.7999 mL/g; CCE-CCK = 0.7620 mL/g), solution density (1.00586 g/mL), and solution viscosity ($1.0206 \times 10-2$ Pa s) were computed using SednTerp. A continuation sedimentation coefficient distribution model of the Lamm Equation, with selection of a maximum entropy model. The molecular weight of each species was calculated using the following equation:

$$c(r) = c_o e^{\frac{M_b \omega^2}{RT}(\frac{r^2 - r_0^2}{2})}$$

Where c(r) is a function of concentration per radius, $c_o$ is the initial concentration, r is the specific radius relative to the reference, $r_o$, ω is the angular velocity, and $M_b$ is the calculated bouyant molecular weight. The quality of fit was determined by the root mean square deviation (RMSD) of the equilibrium data and fit and global score chi-square (GBCS)

**Chemical Synthesis:**

**General.** 1H NMR, 13C NMR, and 19F NMR spectra were recorded on an Agilent 500 MHz VNMRS spectrometer in the stated solvents using tetramethylsilane as the internal standard. Chemical shifts were reported as parts per million (ppm) on the δ scale from the instrument standard. All materials were obtained from Sigma-Aldrich, and unless otherwise stated, were used without further purification. All solvents were purchased as ACS reagent grade or anhydrous if indicated. All reactions were carried out under a nitrogen atmosphere, unless otherwise indicated.

**Preparation of DBCO-Val-Cit-PABC-MMAE and DBCO-Glu-Val-Cit-PABC-MMAE:** Synthesis of both linkers was performed as previously reported by Anami et al.[50] Briefly, chlorotrityl chloride resin (1 eq) was mixed with Fluorenylmethyloxycarbonyl (Fmoc)-citrulline-PABOH (3 eq) in pyridine (3 eq), in 10:1 tetrahydrofuran to dimethylformamide, and agitated overnight at 55°C. After cooling, methanol was added and agitated for 30 min at room temperature. The solution was drained and resin was washed with dimethylformamide and dichloromethane. Fmoc was deprotected after each coupling step with piperidine in dimethylformamide (20%) and washed with dimethylformamide and dichloromethane. Fmoc-protected valine (4 eq) was pre-activated by being mixed with 1-[bis(dimethylamino)methylene]-1H-1,2,3-triazolo[4,5-b]pyridinium 3-oxid hexafluorophosphate (4 eq.) and N,N-diisopropylethylamine (6 equiv.) in dimethylformamide for 2-5 min, followed by mixing with citrulline-PABOH. For the DBCO-Glu-Val-Cit-PABC-MMAE, the washing step and cleavage of Fmoc was repeated, followed by the activation of Fmoc and tert-butyl co-protected glutamic acid.

Following peptide elongation and Fmoc deprotection, the resin was treated with DBCO-$PEG_4$-NHS (2 eq, Broadpharm), 1-[bis(dimethylamino)methylene]-1H-1,2,3-triazolo[4,5-b]pyridinium 3-oxid hexafluorophosphate (2 eq.) and N,N-diisopropylethylamine (3 equiv.) in dimethylformamide for 1 hour, then washed with dimethylformamide and dichloromethane. Addition of MMAE was completed by activation with bis(2,4-dinitrophenyl)carbonate (5 eq) and 4-dimethylaminopyridine (10 eq), followed by addition of MMAE (2.0 eq, MedChem Express) in DMF.

Cleavage of the tert-butyl protecting group on glutamic acid was achieved by addition of trifluoroacetic acid in dichloromethane (20%) to the DBCO-Glu(*t*Bu)-Val-Vit-PABC-MMAE linker for 4 hours at 0°C, followed by quenching with $NH_4OH$.

**CCE-AF488:** AlexaFluor 488 DBCO (MedChemExpress) (2.0 eq) was reacted with CCE peptide in phosphate buffered solution (pH = 7.4) at room temperature for 4 hours while stirring. Following completion of the reaction, the remaining AlexaFluor 488 DBCO was removed with an Amicon 3kDa MWCO spin column (Millipore Sigma), with three sequential washes at room temperature. The retentate was collected and analyzed via NanoDrop Ultra UV-VIS and RP-HPLC.

**CCE-AF647:** AlexaFluor 647 DBCO (MedChemExpress) (2.0 eq) was reacted with CCE peptide in phosphate buffered solution (pH = 7.4) at room temperature for 4 hours while stirring. Following completion of the reaction, the remaining AlexaFluor 647 DBCO was removed with an Amicon 3kDa MWCO spin column (Millipore Sigma), with three sequential washes at room temperature. The retentate was collected and analyzed via NanoDrop Ultra UV-VIS and RP-HPLC.

**CCE-Val-Cit-PABC-MMAE:** Synthesized DBCO-Val-Cit-PABC-MMAE (2.0 eq) was reacted with CCE peptide in phosphate buffered solution (pH = 7.4) at room temperature for 4 hours while stirring. Following completion of the reaction, the remaining DBCO-Val-Cit-PABC-MMAE was removed with an Amicon 3kDa MWCO spin column (Millipore Sigma), with three sequential washes at room temperature. The retentate was collected and analyzed via NanoDrop Ultra UV-VIS and RP-HPLC.

**CCE-Glu-Val-Cit-PABC-MMAE:** Synthesized DBCO-Glu-Val-Cit-PABC-MMAE (2.0 eq) was reacted with CCE peptide in phosphate buffered solution (pH = 7.4) at room temperature for 4 hours while stirring. Following completion of the reaction, the remaining DBCO-Glu-Val-Cit-PABC-MMAE was removed with an Amicon 3kDa MWCO spin column (Millipore Sigma), with three sequential washes at room temperature. The retentate was collected and analyzed via NanoDrop™ One Spectrophotometer UV-VIS and RP-HPLC.

**Fixed Cell Fluorescent Microscopy:** Immunofluorescence staining was performed as previously described.[11] Cells and were labeled with 10 µg/ml α-DEspR$_{SMA}$-568$_2$ for antibody internalization, or treated with 5 µg/ml of α-DEspR$_{SMA}$-MMAE$_2$ and for 5 µg/ml of α-DEspR$_{SMA}$-568$_2$ cytotoxicity experiments. Cells were allowed to bind to 15 minutes at 4°C, followed by removal of media, washing thrice with PBS, and then addition of pre-warmed media. For internalization experiments, cells were treated with lysotracker green (ThermoFisher). For microtubule inhibition, cells were fixed with 2% paraformaldehyde, permeabilized with Triton-X, and then stained with anti-tubulin-AF488 (ThermoFisher). Cells were counter stained with NucBlue Hoechst stain (ThermoFisher). Excitation was performed on 405 nm, 488 nm, and 543 nm laser lines, emission was collected from 430-475 nm, 505-550 nm, and 615-700 nm to prevent overlap. For murine antibody internalization, imaging was performed with a Zeiss Axiostop fluorescence microscope, as previously described127 and confirmed with confocal studies using the Leica SP5, with 1 AU pinhole aperture, 0.7 µm Z-stacks taken for comparison. For antibody internalization, excitation was performed with 405 nm and 543 nm laser lines, emission was collected from 430-475 nm and 600-700 nm to prevent overlap. Quantification was performed using ImageJ with JACoP plugin-software.

**ADC Quantification:**

**Total ADC:** Total ADC was quantified using sandwich ELISA, based on modifications of previously established methods.[2] 96-well Nunc-Immuno MaxiSorp plates (Sigma Aldrich) were coated with a mouse anti-MMAE antibody (4 μg/mL) [BioRad] in bicarbonate buffer and incubated overnight at 4 °C. After washing with 0.05% Tween in PBS (PBS-T) thrice and blocking with 1% BSA for 4 hrs at 4°C, ADC standard with serial dilutions and samples were added, and incubated at 37 °C for 1 h. Quality control samples were prepared for the assay to ensure adequate sample detection, similar to previous studies.[52] After samples were then washed thrice with 0.1% PBS-T, 100 μL of an HRP-conjugated goat anti-mouse IgG2b [Invitrogen] solution (100 ng/mL) was added and the plate was incubated at 37 °C for 1 h. After washing the plate thrice with 0.1% PBS-T, TMB (BioRad) substrate was added and the reaction was stopped with 2 M $H_2SO4$ before detection of the absorbance signal at 450 nm on a SpectraMax 7 Plate Reader.

**Total Antibody:** Total antibody was quantified by sandwich ELISA, based on modifications of previously established methods.[52] 96-well Nunc-Immuno MaxiSorp plates (Sigma Aldrich) were coated with 10 μg/ml of the antigen peptide ($M_1$TMFKGSN$E_9$) in bicarbonate buffer, and incubated overnight at 4°C. Plates were washed with phosphate buffered saline (PBS) thrice, followed by blocking with 200 μl 1% bovine serum albumin (BSA) in PBS for 4 hours at 4°C. After washing thrice with PBS, antibody standard with serial dilutions and samples were added, and incubated at 37 °C for 1 h. Quality control samples were prepared for the assay to ensure adequate sample detection, similar to previous studies.[52] Samples were then washed thrice with 0.1% After samples were then washed thrice with 0.1% PBS-T, 100 μL of an HRP-conjugated goat anti-mouse IgG2b [Invitrogen] solution (100 ng/mL) was added and the plate was incubated at 37 °C for 1 h. After washing the plate thrice with 0.1% PBS-T, TMB (BioRad) substrate was added and the reaction was stopped with 2 M $H_2SO4$ before detection of the absorbance signal at 450 nm on a SpectraMax 7 Plate Reader.

**Fluorescence Anisotropy Experiments:** Studies were performed on a SpectraMax 7 Plate Reader, using fluorescence and fluorescence anisotropy settings. Samples were run using black wall 96 well plates (Corning). Initial fluorescence readings were performed from titrations at 300 nM to 1 nM concentrations in 100 μl in triplicate. Anisotropy was calculated for samples over this range, using the following formula:

$$r = \frac{I_{vv} - GI_{VH}}{I_{vv} + 2GI_{VH}}$$

With G calculated from the individual AF647 fluorophore. Background subtraction was performed for each sample. A concentration of 3 nM was selected for quantification of CCE-AF647 and α-DEsp$R_{SMA}$-$647_2$ based on the appropriate signal to noise ratio. Samples were prepared in varying concentrations of albumin, and experiments were performed after 24 hours of mixing at 37°C. Samples contained either 1% DMSO or no DMSO to evaluate for interaction between hydrophobic binding pockets.

**Anti-IgM Quantification:** Total anti-drug IgM antibodies were detected using previously reported methods.[53] 96-well Nunc-Immuno MaxiSorp plates (Sigma Aldrich) were coated with α-DEsp$R_{SMA}$-MMA$E_2$ (4 μg/mL) or α-DEsp$R_{SMA}$ mAb and incubated overnight at 4 °C. After washing with 0.1% PBS-T thrice and blocking with 1% BSA for 4 hours at 4°C, anti-IgG2b IgM rat (Novus Biologics) standard with serial dilutions or serum samples and incubated at 37 °C for 1 h. After washing, 100 μL of an HRP-conjugated goat anti-rat IgM was added and the plate was incubated at 37 °C for 1 h. After washing the plate thrice with 0.1% PBST, TMB substrate was added and the reaction was stopped with 2 M H2SO4 before detection of the absorbance signal at 450 nm.

**Anti-IgG Quantification:** Total anti-drug IgM antibodies were detected using previously reported methods.[53] 96-well Nunc-Immuno MaxiSorp plates (Sigma Aldrich) were coated with α-DEspR$_{SMA}$-MMAE$_2$ (4 µg/mL) or α-DEspR$_{SMA}$ mAb and incubated overnight at 4 °C. After washing with 0.1% PBST thrice and blocking with 3% BSA, anti-IgG2b IgG rat (Novus Biologics) standard with serial dilutions or serum samples and incubated at 37 °C for 1 h. After washing, 100 µL of an HRP-conjugated goat anti-rat IgG was added and the plate was incubated at 37 °C for 1 h. After washing the plate thrice with 0.1% PBST, TMB substrate was added and the reaction was stopped with 2 M H2SO4 before detection of the absorbance signal at 450 nm.

| Conjugation Method | Site Selective | Uniform Drug Loading | Glycosylation-Independent | Versatile Vector Selection |
|---|---|---|---|---|
| **Modified cysteine residues (ex: thiomab)** | Yes | No | Yes | Yes |
| **Modified peptide sequence (ex: π-clamp)** | Yes | No | Yes | Yes |
| **Unnatural amino acids (ex: pAMF)** | Yes | No | No | No |
| **Enzymatic Conjugation (ex: transglutaminase)** | Yes | No | No | No |
| **Peptide Tags (ex: transglutaminase)** | Yes | No | No | Yes |

**Table S1. Comparison of current landscape of site-specific conjugation.**

Contemporary methods of site-specific conjugation are shown, with their ability to enable site-specific modification, uniform (100% conversion of ADC at all conjugation sites) drug loading, requirement of altered glycosylation of the antibody (glycosylation-dependent changes) to allow conjugation, and whether vector selection is not a factor in conjugation.

| Peptide ID | Peptide Type | Sequence |
|---|---|---|
| Peptide L/E | Acidic | MKLEEILSELEEILSELEEILYELEEILSEVGER |
| Peptide L/K | Basic | MKLKKIKSLLKKIKSLLKKIKSLLKKIKSLVGER |
| Peptide V/E | Acidic | MKLEEIVSELEEIVSELEEIVYELEEIVSEVGER |
| Peptide V/K | Basic | MKLKKIKSVLKKIKSVLKKIKSVLKKIKSVVGER |
| Peptide I/E | Acidic | MKLEEIISELEEIISELEEIIYELEEIISEVGER |
| Peptide I/K | Basic | MKLKKIKSILKKIKSILKKIKSILKKIKSIVGER |

**Table S2.** Sequences of modified leucine-zipper sequences

Prepared sequences of each peptide, labeled for the *g*/*e'* site that was modified by a hydrophobic amino acid, followed by whether glutamic acid (E) or lysine (K) was used.

| | RMSD of Fit | | | |
|---|---|---|---|---|
| Species | Single | Monomer/Dimer | Monomer/Trimer | Monomer/Tetramer |
| CCE | **0.005442** | 0.005684 | 0.005684 | 0.005684 |
| CCK | **0.005281** | 0.005422 | 0.005682 | 0.005422 |
| CCE-CCK | 0.005791 | 0.01106 | 0.01836 | **0.004801** |

**Table S3.** Quality of Model Fit for Each Oligomeric State.

| Species | Best Predictive Model | RMSD of Fit | $K_{d,\ tetramer}$ |
|---|---|---|---|
| CCE | Single Species | 0.005442 | 0.995 M |
| CCK | Single Species | 0.005281 | 0.0971 M |
| CCE-CCK | Monomer → Tetramer | 0.004801 | $1.1021 \times 10^{-10}$ M |

**Table S4.** Best predictive fit of each species and estimated dissociation constant

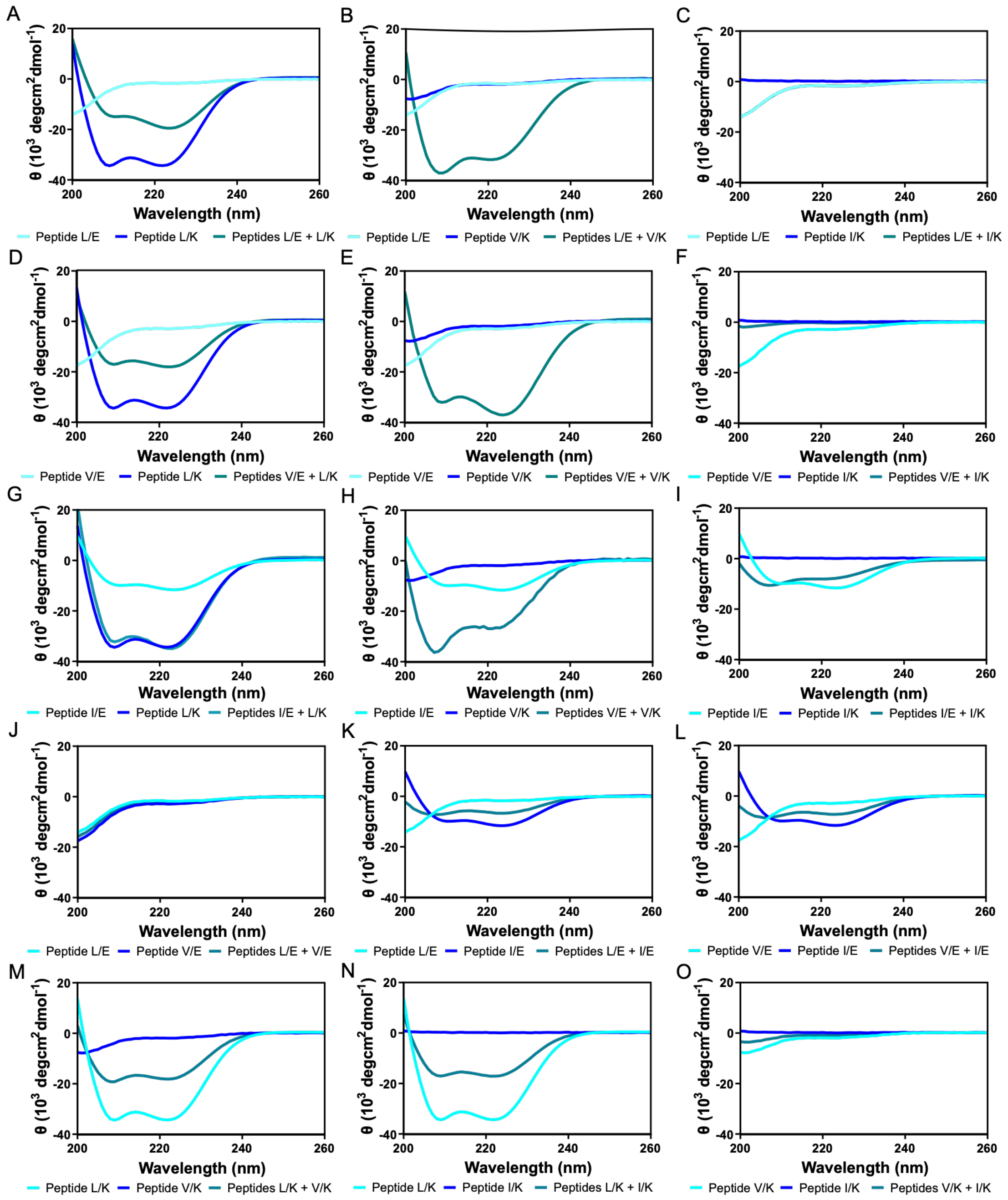


**Fig. S1. (A-I).** Circular dichroism spectrum of all peptide acidic and basic pairings. Peptides are labeled by the hydrophobic amino acid in either the *e'* or *g* (first letter; L is leucine, V is valine, and I is isoleucine), and whether they are basic (second letter, K) or acidic (second letter E). **(J-**

**L).** Circular dichroism of different acidic peptide pairings, demonstrating that mixtures of **(J)** L/E (red) and V/E (blue), **(K)** L/E (red) and I/E (blue), or **(L)** V/E (red) and I/E (blue) do not form supramolecular structures, likely because of charge repulsion between the species. **(M-O).** Circular dichroism of basic peptide pairings, demonstrating that mixtures of **(M)** L/K (red) and V/K (blue), **(N)** L/K (red) and I/K (blue), or **(O)** V/K (red) and I/K (blue) do not form supramolecular structures, as equimolar spectra (purple) are an average of the two species.

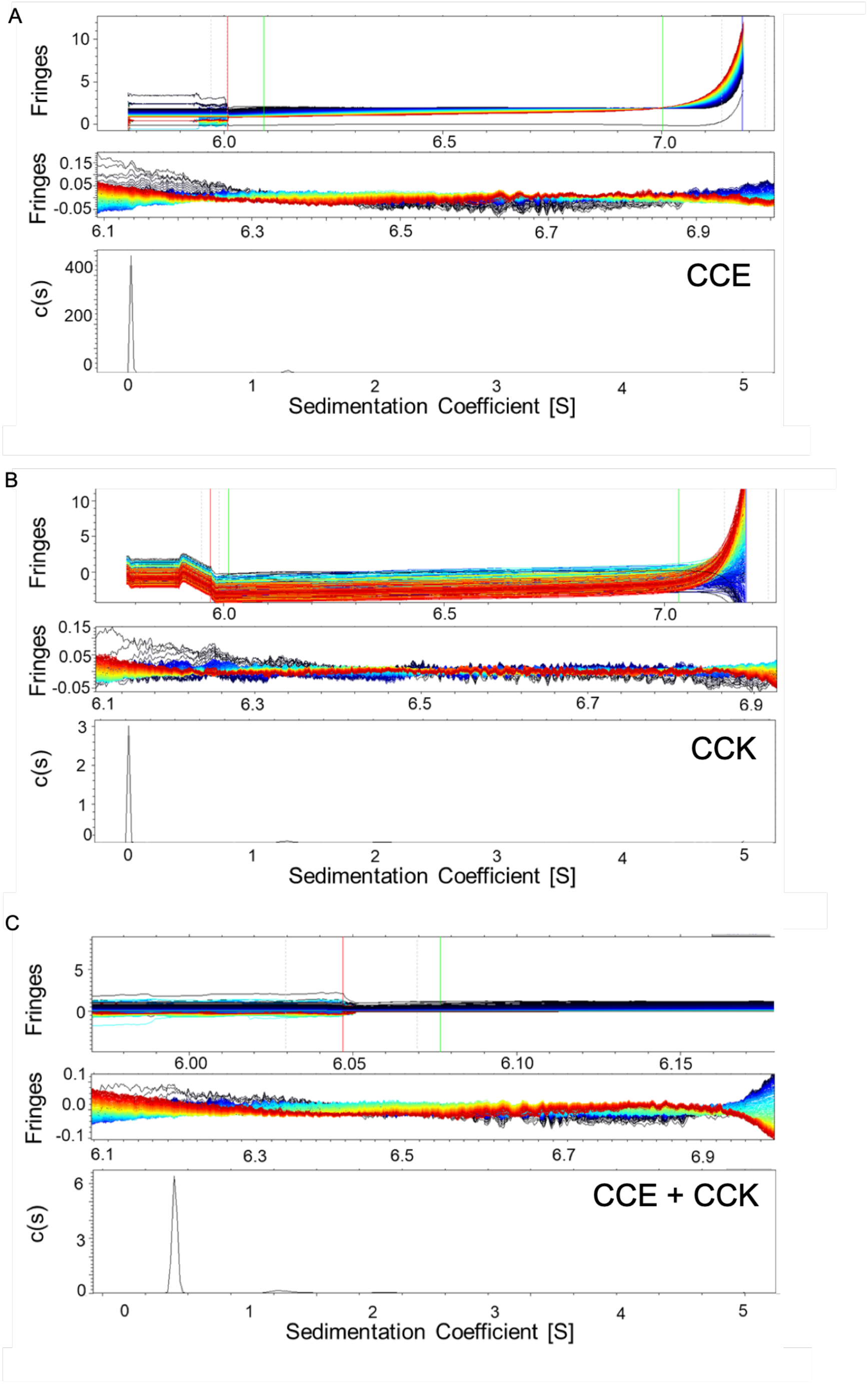


**Fig. S2. (A-C)** Analytical ultracentrifugation velocity scans, performed with 0.6 mg/mL of total peptide, using interference optics, were performed to analyze the molecular weight of CCE, CCK, and a CCE-CCK mixture, providing information of the formation of oligomeric states. **(A)** Velocity scans of peptide CCE, at demonstrating a single species with a low molecular weight. Using the modified Svedberg equation, this gives a molecular weight of approximately 4,000 kDa. **(B)** Velocity scans of peptide CCK, also demonstrating a single species with a low molecular weight. Using the modified Svedberg equation, this gives a molecular weight of approximately 4,000 kDa. **(C)** Velocity scans of a 1:1 mixture of peptide CCE-CCK, which again demonstrates a single predominant species; however, there is a shift in molecular weight. Using the modified Svedberg equation, this provides a calculated weight of 16,281 kDa (theoretical 15,788 kDa), supporting heterotetramer formation.

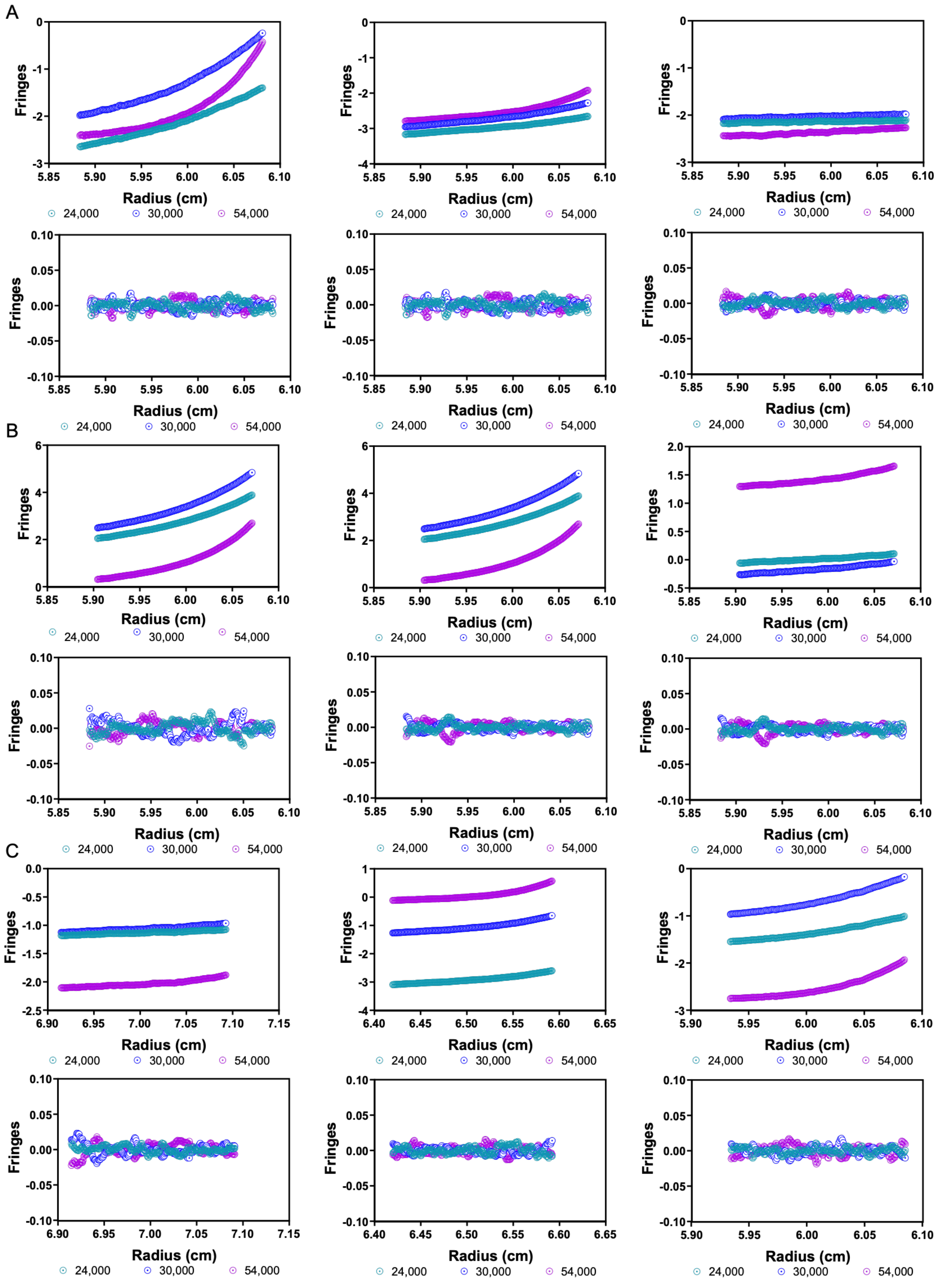


**Fig. S3. (A-C)** Analytical ultracentrifugation scans were performed using interference optics, with varying concentrations (0.2 mg/mL, left; 0.6 mg/mL, middle; 1.0 mg/mL, right) and varying rotational speeds (24,000 rpm, teal; 30,000 rpm, blue; and 54,000 rpm, purple), to provide more precise information on oligomeric species formation and strength of interaction between peptides. **(A)** Equilibrium scans of peptide CCK (top) with residuals (bottom), demonstrating a buoyant molecular weight ($M_B$) of 4,288 kDa, with weak association ($K_d$ = 0.0971 M), and fitting a single species model with good fidelity (RMSD of fit: 0.005281). **(B)** Equilibrium scans of peptide CCE (top) with residuals (bottom), demonstrating a buoyant molecular weight ($M_B$) of 4,241 kDa, with weak association ($K_d$ = 0.995 M), and fitting a single species model with good fidelity (RMSD of fit: 0.005442). **(C)** Equilibrium scans of equimolar peptide CCE-CCK (top) with residuals (bottom), demonstrating a buoyant molecular weight ($M_B$) of 15,306 kDA, with strong association ($K_d = 1.125 \times 10^{-10}$), and fitting a tetrameric association model with good fidelity (RMSD of fit: 0.004801)

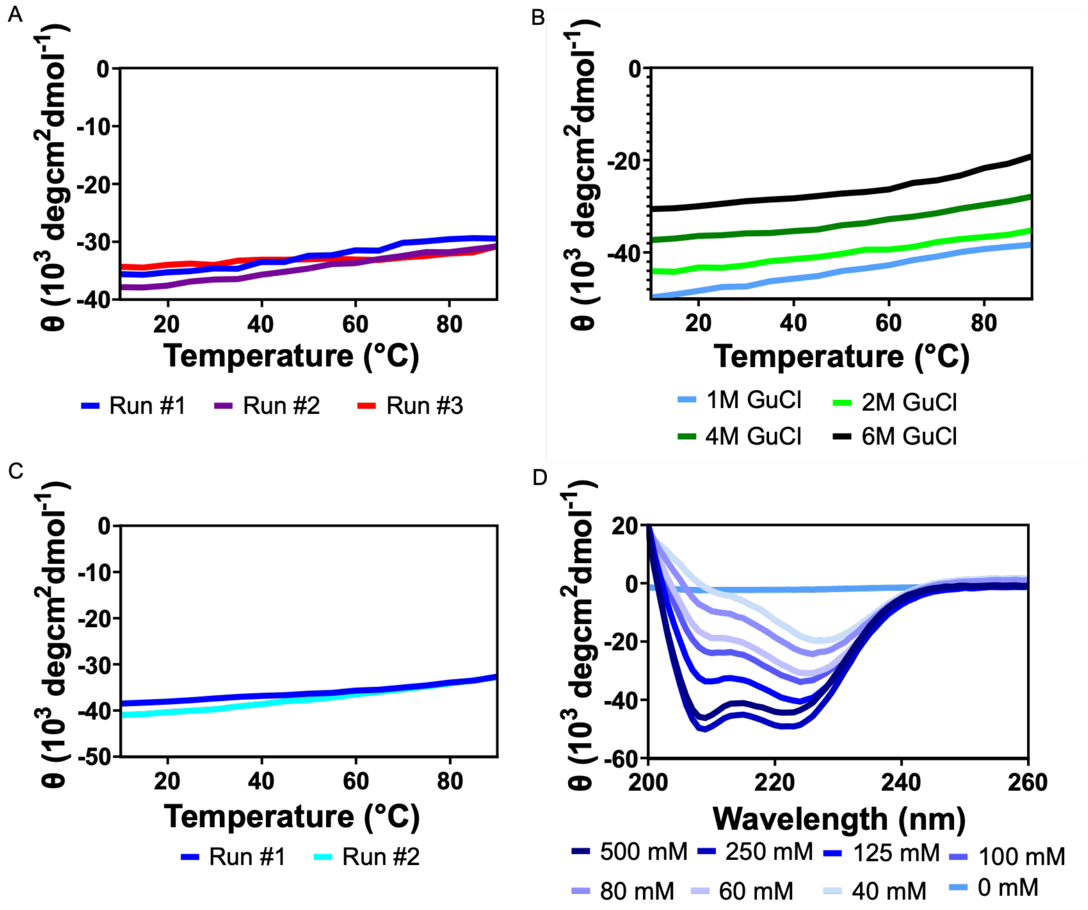


**Fig. S4. (A)** Thermal denaturing study of CCE-CCK, with representative spectrum shifts at 222 nM, ranging from 10°C to 90°C, which reflects the shift in the amide backbone and demonstrates the degree of retained coiled structure. At 90 ˚C, only 18.2% + 0.8% of the CCE-CCK construct unfolds. The repeat runs (blue, purple, and red), reflect the reversibility of this folding, demonstrating that aggregation does not form. **(B)** Chaotropic and thermal denaturing study of CCE-CCK, with representative spectra at 222 nm of samples equilibrated in 1M (light blue), 2M (light green), 4M (dark green), and 6M (black) guanidium chloride (GuCl), as temperature increases from 10°C to 90°C. Protein folding is observed, up to 53.5% at 90°C in 6M guanidinium chloride (GuCl). **(C)** Circular dichroism of peptides CCE-CCK in equimolar ratio at pH = 3, run with repeat thermal melting from 5ºC to 90ºC. While thermal melting was observed, this was reversible with cooling. **(D)** The impact of salinity was assessed by dialyzing the coil structure into different buffered solutions. As the salt concentration (measured as mM of sodium chloride) was decreased, the structure became less stable, ultimately collapsing without the presence of salt (dialysis of sample in deionized water).

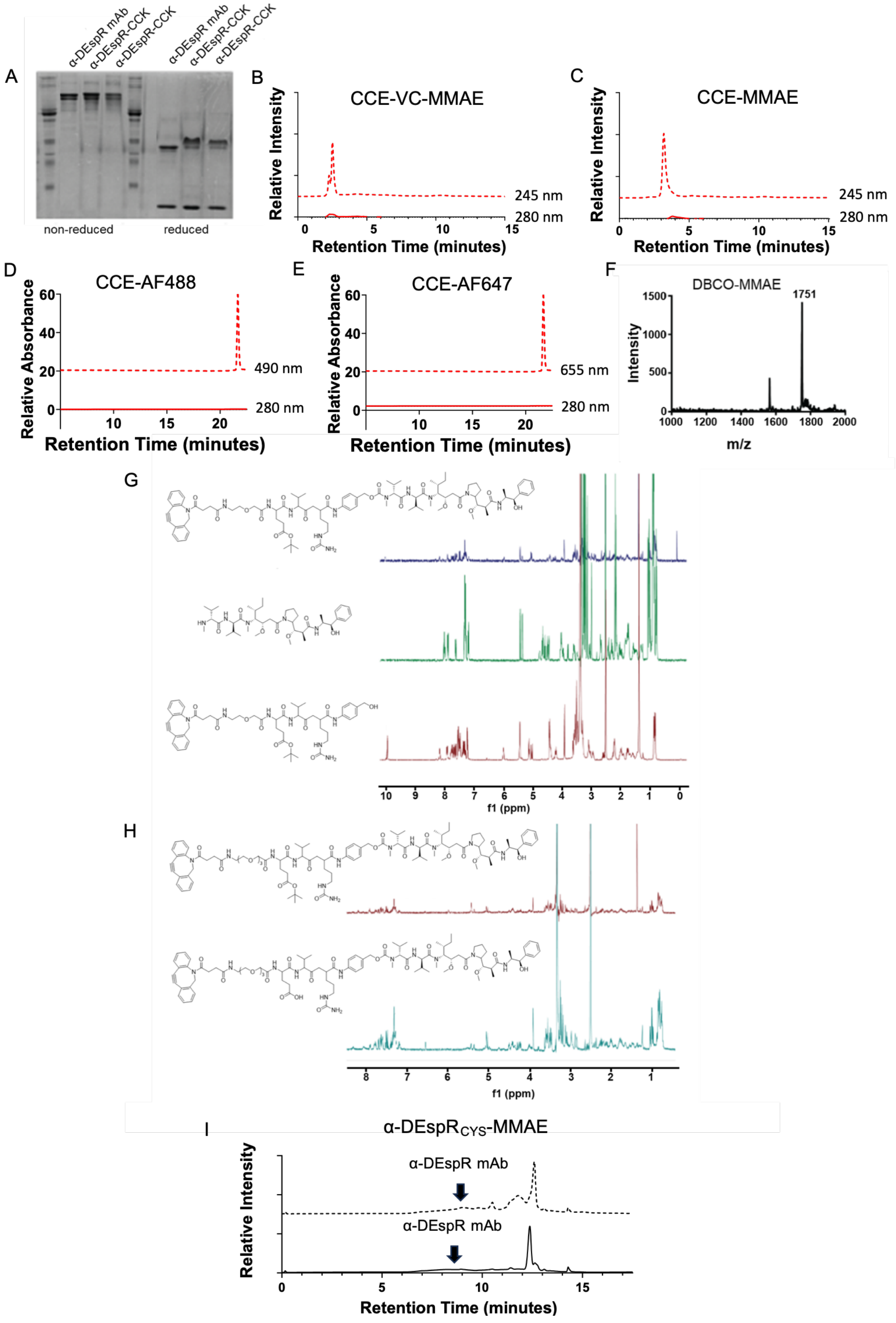


**Fig. S5. (A)** SDS-PAGE (left non-reduced, right reduced), showing the difference in modified α-DEspR-CCK from α-DEspR antibody. **(B)** Representative chromatogram showing the modified CCE-vc-MMAE peptide after drug loading. **(C)** Representative chromatogram showing the modified CCE-EVC-MMAE peptide after drug loading. **(D)** Representative chromatogram showing the modified CCE-AF-488 peptide after drug loading. **(E)** Representative chromatogram showing the modified CCE-AF-647 peptide after drug loading. **(F)** LC-MS tracing confirming the molecular weight of the DBCO-EVC-MMAE linker. **(G)** $^{13}C$ NMR spectra showing (top) the DBCO-EVC-MMAE linker (with glutamic acid protection), (middle) MMAE, and (bottom), the DBCO-EVC-PABC structure prior to addition of MMAE. **(H)** $^{1}H$ NMR showing (top) the DBCO-EVC-MMAE linker (with glutamic acid protection) and (bottom) the DBCO-EVC-MMAE linker after deprotection. **(I)** Representative chromatogram showing the cysteine conjugated α-DEspR$_{CYS}$-MMAE ADC, with peak for unconjugated mAb highlighted.

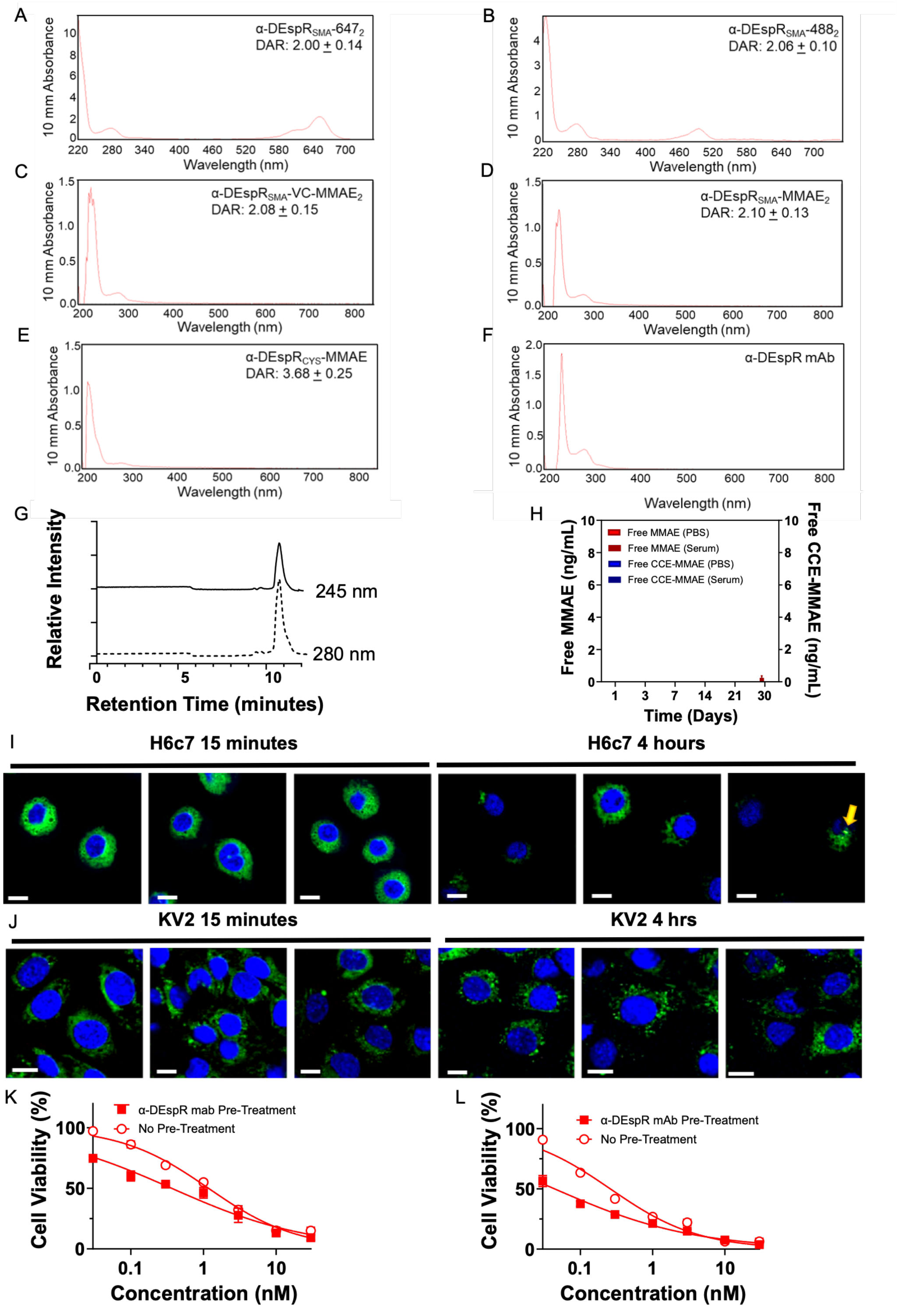


**Fig. S6. (A)** Representative UV-VIS spectrum of α-DEspR$_{SMA}$647$_2$ (DAR=2.00±0.14). **(B)** Representative UV-VIS spectrum α-DEspR$_{SMA}$488$_2$ (DAR=2.06±0.10). **(C)** Representative UV-VIS spectrum of α-DEspR$_{SMA}$-VC-MMAE$_2$ (DAR=2.08±0.15). **(D)** Representative UV-VIS spectrum of α-DEspR$_{SMA}$-MMAE$_2$ (DAR=2.10±0.13). **(E)** Representative UV-VIS spectrum of α-DEspR$_{CYS}$MMAE (DAR=3.68±0.25). **(F)** Representative UV-VIS of α-DEspR mAb. **(G)** Representative RP-HPLC chromatogram of α-DEspR$_{SMA}$ -VC-MMAE$_2$. **(H)** *Ex-vivo* stability assay of α-DEspR$_{SMA}$-MMAE$_2$, evaluating free MMAE in PBS (red) or rat serum (dark red) or CCE-MMAE release in PBS (blue) or rat serum (dark blue). No free drug and minimal free drug (<1%) were detected after 30 days. Free peptide was not detected in the samples using RP-HPLC. **(I,J)** Representative confocal microscopy of α-DEspR$_{SMA}$647$_2$ (red) internalization in **(I)** DEspR-low H6c7 and **(J)** DEspR-negative KV2 cells at 15 minutes vs 4 hours. Cells were stained with NucBlue (blue, nucleus) and Lysotracker (green, lysosome). Minimal α-DEspR$_{SMA}$647$_2$ uptake was noted in H6c7 cells and no uptake was noted in KV2 cells. **(K,L)** MMAE cytotoxicty at 24 hours were compared either with (red box) or without (hollow circle) 24hrs of α-DEspR mAb pre-treatment. Pre-treatment with α-DEspR mAb improved cytotoxicity of MMAE in **K)** Panc1 ($IC_{50,24h}$: 0.365±0.04 vs 1.173±0.07 nM) and **L)** MiaPaCa2 cells ($IC_{50,24h}$: 0.044±0.003 vs 0.260±0.020 nM).

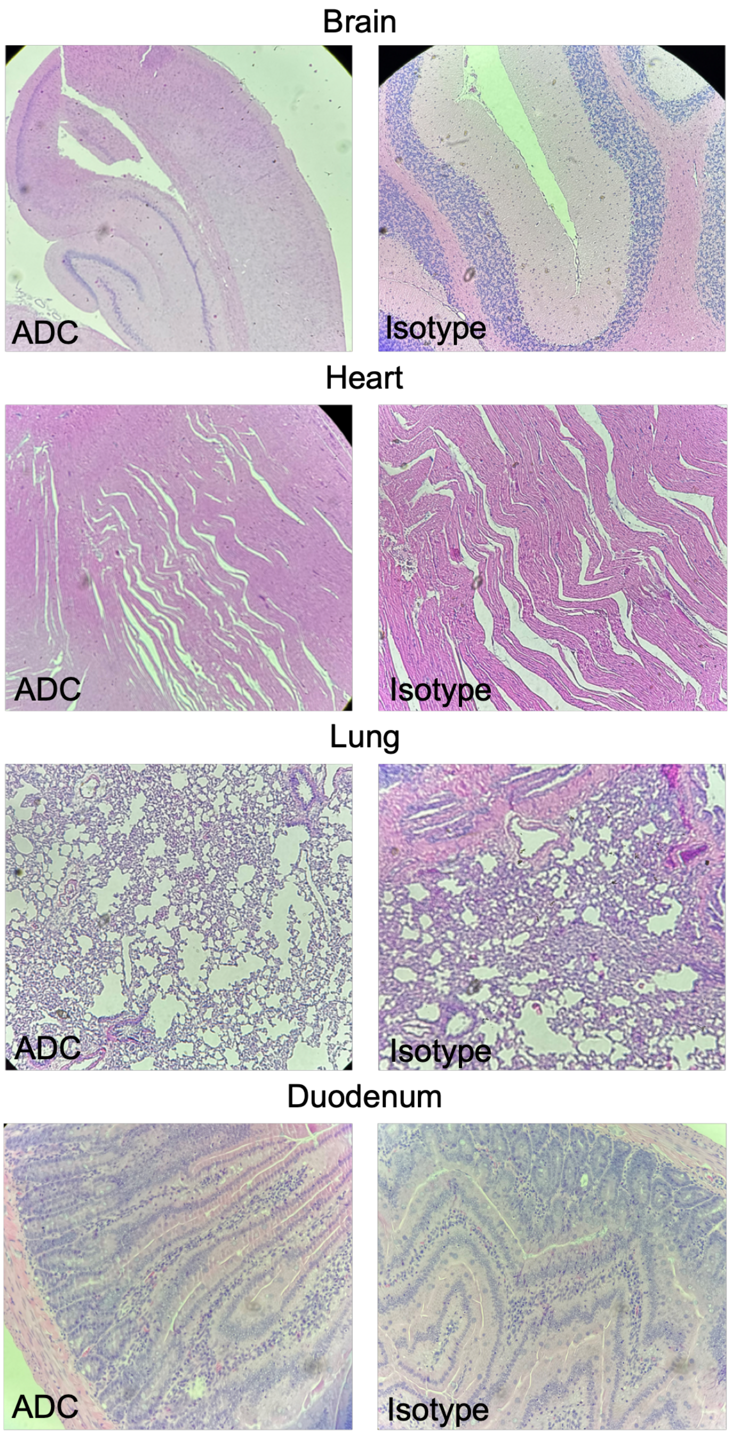


**Fig. S7.** Representative H&E stains of rat organs for biodistribution study.

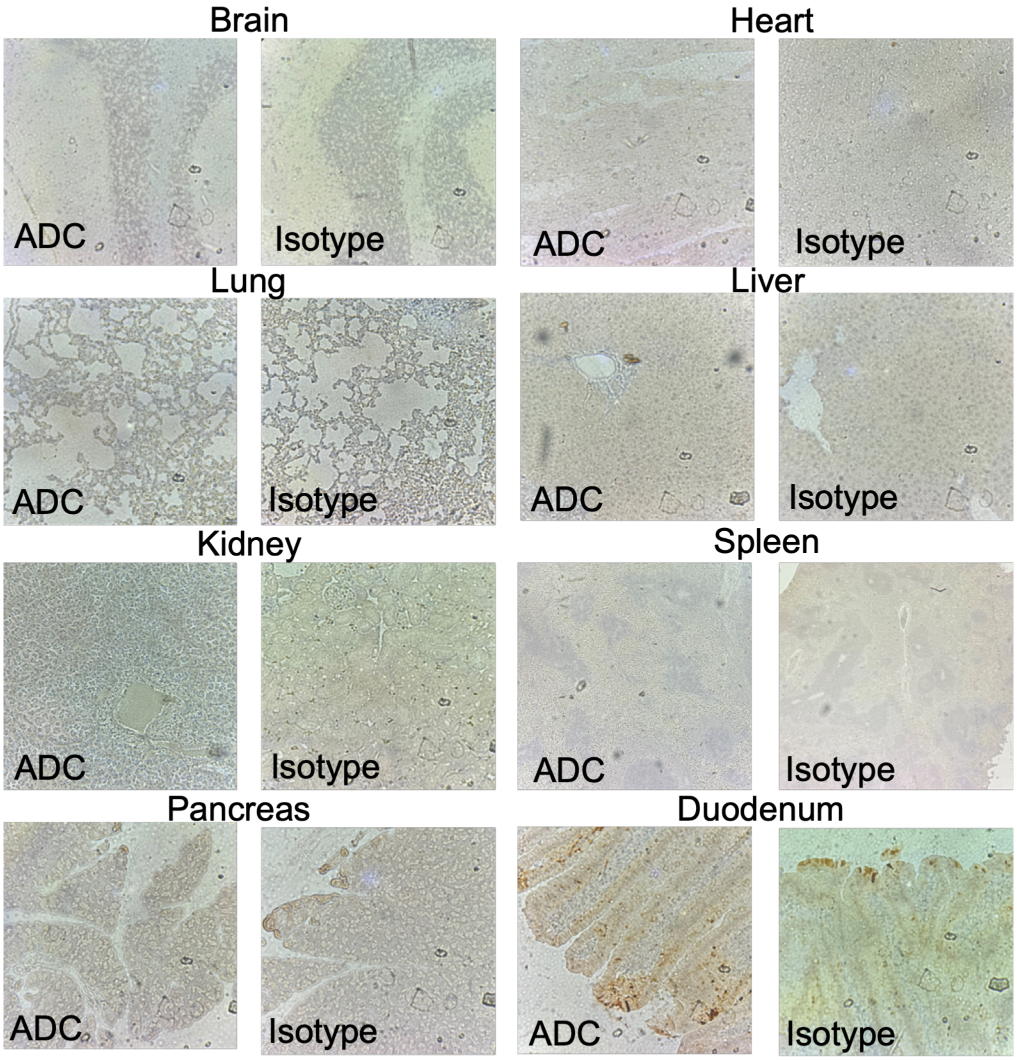


**Fig. S8.** Representative Caspase-3 DAB staining of biodistribution organs

A

| Treatment Group | GOO | Perforation | Intestinal Ischemia | Porta Hepatis Infiltration |
|---|---|---|---|---|
| Saline | 4 | 0 | 2 | 2 |
| α-DEspR mAb | 3 | 0 | 2 | 3 |
| α-DEspR$_{CYS}$-MMAE | 5 | 1 | 1 | 1 |
| α-DEspR$_{SMA}$-MMAE$_2$ | 0 | 0 | 0 | 0 |
| Total | 12 | 1 | 5 | 6 |

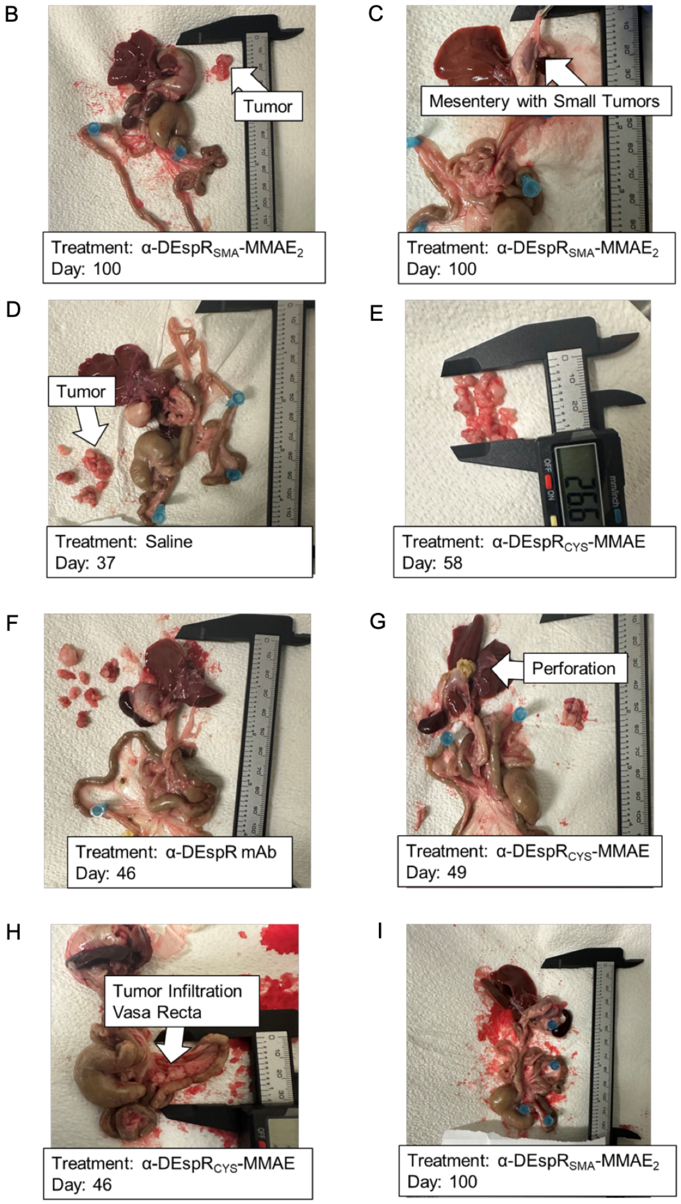


**(A)** Clinical data of rats at time of euthanasia, with cause of death if euthanasia was required prior to 100 days. Common etiologies included gastric outlet obstruction (GOO), defined as visible evidence of tumor invasion into the proximal duodenum or antrum/pylorus of the stomach, perforation of gastrointestinal viscus, visualized with initial laparotomy, intestinal ischemia, defined as visible evidence of necrotic bowel with tumor invasion into adjacent vasculature, or persistent jaundice with evidence of invasion into biliary system. **(B-I)** Representative pictures of rats at time of euthanasia. (**B,C)** In α-DEspR$_{SMA}$-MMAE$_2$ treated rats with visible tumor, there were either **(B)** few solitary tumors **(C)** or small, hard to characterize tumors within the peripancreatic mesentery, confirmed with histologic staining. Tumor burden in **(D)** saline rats was comparable to **(E)** α-DEspR$_{CYS}$-MMAE (tumor only) and **(F)** α-DEspR mAb treated rats. **(G)** Perforation with leakage of gastric contents noted at time of euthanasia. Salvaged bowel demonstrated a leak near a site of tumor invasion. **(H)** Tumor invasion into mesenteric vessels, leading to progressive weight loss and intestinal ischemia. **I)** Representative organs from 1/3 α-DEspR$_{SMA}$-MMAE$_2$ without visualized tumor at the time of euthanasia. Tumor was noted within the peri-pancreatic mesentery.
